# The *Legionella* Lqs-LvbR regulatory network controls temperature-dependent growth onset and bacterial cell density

**DOI:** 10.1101/2021.10.12.464170

**Authors:** Ramon Hochstrasser, Hubert Hilbi

## Abstract

*Legionella* species are facultative intracellular pathogens, which can cause a life-threatening pneumonia termed Legionnaires’ disease. *Legionella pneumophila* employs the *Legionella* quorum sensing (Lqs)-LvbR network to regulate virulence and motility, but its role for growth in media is ill-defined. Compared to the parental *L. pneumophila* strain JR32, a Δ*lqsR* mutant showed a reduced lag phase at 30°C and reached a higher cell density at 45°C, while the Δ*lqsA*, Δ*lqsS* and Δ*lqsT* mutants exhibited a longer lag phase and reached only a lower cell density. A Δ*lvbR* mutant resumed growth like the parental strain at 30°C, but exhibited a substantially reduced cell density at 45°C. Thus, LvbR is an important cell density regulator at elevated temperatures. A quantitative analysis of temperature-dependent growth characteristics of environmental and clinical strains revealed that *L. pneumophila* strains grew in AYE medium after distinct lag phases with similar rates at 30°C, reached different cell densities at the optimal growth temperature of 40°C, and no longer grew at 50°C. *Legionella longbeachae* reached a rather low cell density at 40°C and did not grow at and beyond 45°C. Genes encoding components of the Lqs-LvbR network were present in the genomes of the environmental and clinical *L. pneumophila* isolates, and the P_*lqsR*_, P_*lqsA*_, P_*lqsS*_ and P_*lvbR*_ promoters from strain JR32 were active in these strains. Taken together, our results indicate that the Lqs-LvbR network governs the temperature-dependent growth onset and cell density of the *L. pneumophila* reference strain JR32, and possibly also of environmental and clinical *L. pneumophila* isolates.

**Importance:** Environmental bacteria of the genus *Legionella* are the causative agents of the severe pneumonia Legionnaires’ disease, the incidence of which is worldwide on the rise. *Legionella pneumophila* and *Legionella longbeachae* are the clinically most relevant species. The opportunistic pathogens are inhaled through contaminated aerosols and replicate in human lung macrophages with a similar mechanism as in their natural hosts, free-living amoebae. Given their prevalence in natural and technical water systems, an efficient control of *Legionella* spp. by physical, chemical or biological means will reduce the incidence of Legionnaires’ disease. Here we report that the *Legionella* quorum sensing (Lqs) system and the pleiotropic transcription factor LvbR govern the temperature-dependent growth onset and cell density of bacterial cultures. Hence, the growth of *L. pneumophila* in water systems is not only determined by the temperature and nutrient availability, but also by quorum sensing, i.e., density- and signaling molecule-dependent gene regulation.

## Introduction

Environmental bacteria of the genus *Legionella* are the causative agents of Legionnaires’ disease (1, 2). This life-threatening pneumonia almost exclusively affects elderly people, can occur in epidemics of several hundreds of cases and is globally on the rise. A total of 65 *Legionella* species has been identified to date (3), of which at least 24 have been linked to human disease (1). However, more than 90% of the Legionnaires’ disease cases diagnosed is caused by *Legionella pneumophila*, which together with *Legionella longbeachae* account for ca. 95% of all *Legionella* infections (1).

*L. pneumophila* parasitizes free-living amoebae and other protozoa (4–6), wherein which it forms a unique replication-permissive compartment, the *Legionella*-containing vacuole (LCV) (7–12). *L. pneumophila* is a facultative intracellular bacterium, which can also grow extracellularly in complex and defined media as well as in biofilms (13, 14). For intra- and extracellular growth, *L. pneumophila* requires micronutrients such as iron (15, 16). The analysis of the carbon and energy sources of *L. pneumophila* revealed that this fastidious, obligate aerobic bacterium mainly metabolizes amino acids (such as serine), but also uses carbohydrates and polysaccharides (reviewed in (17)). Accordingly, *L. pneumophila* utilizes glucose (18, 19) and inositol (20), or glycerol (21) as well as glycerolipids (22), and possibly polysaccharides (glycogen, starch, cellulose, chitin) (23). In general, *L. pneumophila* employs a bi-partite metabolism (24), as amino acids (e.g., serine) are preferentially used catabolically as energy supply, and carbohydrates (e.g., glucose) as well as glycerol are primarily utilized for anabolic processes.

*L. pneumophila* employs a bi-phasic life cycle, where growth and virulence are reciprocally linked: growing bacteria are non-virulent, while non-growing bacteria are virulent, motile and stress resistant (transmissive) (17, 25). The bi-phasic life cycle is controlled by the RNA-binding, post-transcriptional global regulator CsrA (carbon storage regulator A), by the two-component systems LetAS, PmrAB and CpxRA (26), as well as by the *Legionella* quorum sensing (Lqs) system (27, 28). The Lqs system promotes bacterial density-dependent gene regulation, and produces, detects and responds to the low molecular weight signaling molecule LAI-1 (*Legionella* autoinducer-1, 3-hydroxypentadecane-4-one) (29, 30). The components of the Lqs system are encoded in a cluster (*lqsA*–*lqsR*–*hdeD*–*lqsS*) (31, 32) and at a distant site in the *L. pneumophila* genome (*lqsT*) (33), and they are all expressed from distinct promoters (34).

The constituents of the Lqs system comprise the pyridoxal-5-phosphate-dependent autoinducer synthase LqsA (35), the membrane-bound homologous sensor histidine kinases LqsS and LqsT (33, 35), and the prototypic response regulator LqsR (36) (**Fig. 1**). The latter contains a canonical phosphate receiver domain, dimerizes upon sensor kinase-mediated phosphorylation, and harbors an output domain structurally related to nucleotide-binding proteins (37, 38). Intriguingly, the Lqs system is linked to c-di-GMP signaling through the pleiotropic transcription factor LvbR, which negatively regulates *hnox1* expression (39, 40), and itself is negatively regulated by LqsS (35, 39) (**Fig. 1**). The Hnox1-Lpg1057 system comprises the diguanylate cyclase inhibitor Hnox1 and the GGDEF/EAL domain-containing diguanylate cyclase Lpg1057 (41, 42).

**FIG 1.**
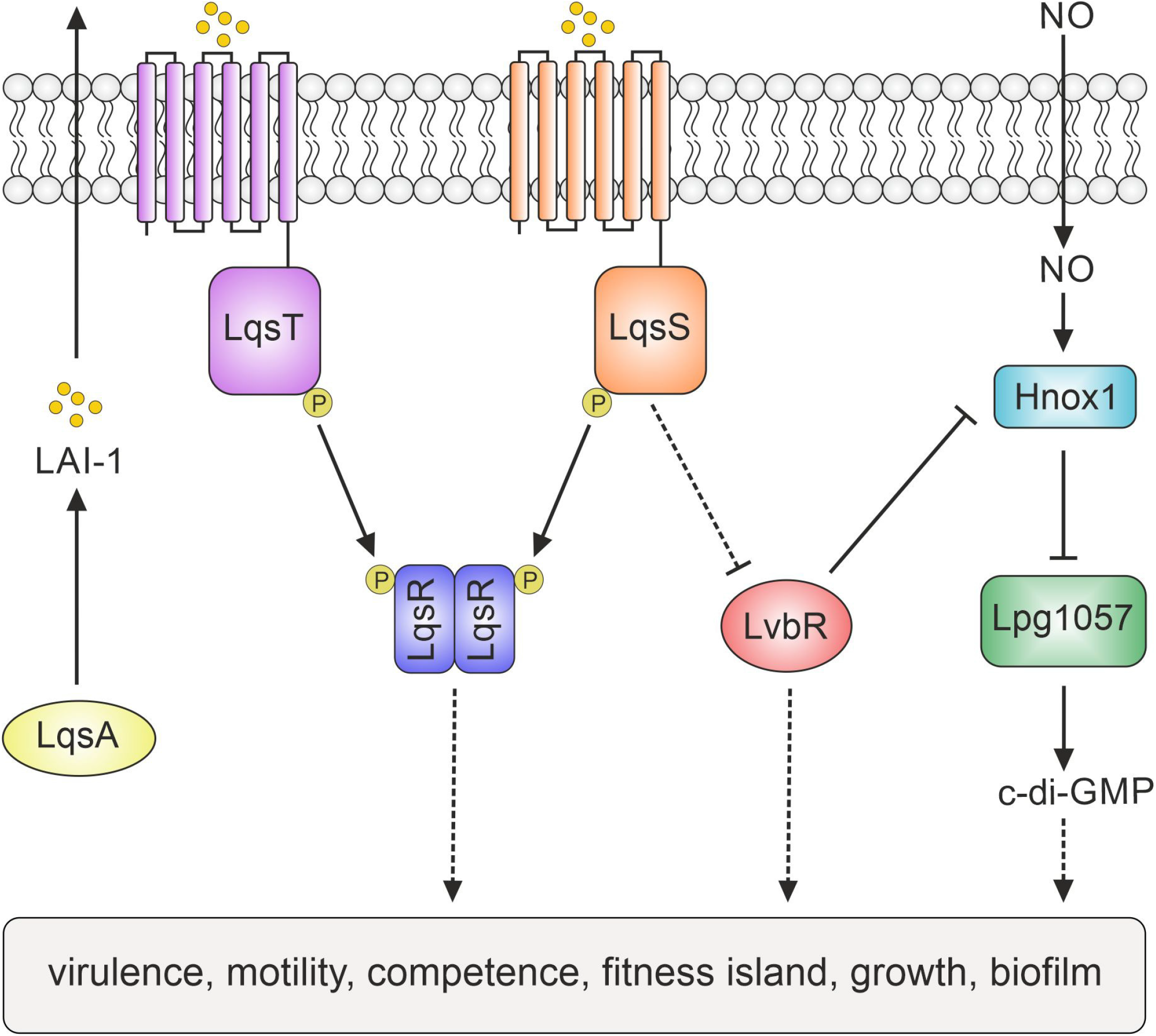
The *Legionella* quorum sensing (Lqs)-LvbR-cyclic di-GMP regulatory network. The environmental bacterium *L. pneumophila* employs a complex regulatory network to integrate and respond to extra- and intracellular signals. The *Legionella* quorum sensing (Lqs) system produces, detects and responds to the endogenous signaling molecule LAI-1 (*Legionella* autoinducer-1; 3-hydroxypentadecane-4-one). The Lqs system comprises the autoinducer synthase LqsA, the homologous sensor kinases LqsS and LqsT, as well as the cognate response regulator LqsR. Lpg1057 is a diguanylate cyclase, which produces the intracellular second messenger cyclic di-GMP (c-di-GMP). Hnox1 is a receptor of nitric oxide (NO) and an inhibitor of the diguanylate cyclase Lpg1057. The transcription factor LvbR negatively regulates *hnox1*expression, and *lvbR* expression is negatively regulated by LqsS. Thus LvbR links the Lqs system and c-di-GMP signaling. The Lqs-LvbR regulatory network controls bacterial virulence, motility, natural competence, expression of a genomic island, planktonic growth and biofilm architecture.

The Lqs-LvbR regulatory network controls pathogen-host cell interactions, bacterial motility, filament production, natural competence for DNA uptake, and biofilm architecture as well as the expression of a 133 kb genomic “fitness island” (39, 40, 43) (**Fig. 1**). The Lqs system and synthetic LAI-1 not only regulate flagellum production and motility of *L. pneumophila* (30), but also the migration of eukaryotic cells, and accordingly, the bacterial signaling compound promotes inter-kingdom signaling (44). Recent studies revealed that the Lqs system also determines phenotypic heterogeneity and the occurrence of functionally different *L. pneumophila* subpopulations in infected amoebae and under sessile/biofilm conditions (45–47). In this work, we show that the Lqs-LvbR system governs the temperature-dependent growth onset and cell density of an *L. pneumophila* reference strain and possibly also of environmental and clinical *L. pneumophila* isolates.

## Results

### The Lqs-LvbR network regulates *L. pneumophila* growth and density in AYE medium

The *L. pneumophila* Lqs-LvbR regulatory network controls a number of bacterial traits, including virulence and motility, natural competence and biofilm architecture (40), but its role for planktonic bacterial growth and temperature dependency of replication has not been studied in detail. To address this question, we compared growth of the parental strain JR32 with the Δ*lqsR*, Δ*lqsA*, Δ*lqsS*, Δ*lqsT*, Δ*lqsS*-Δ*lqsT* or Δ*lvbR* mutant strains in AYE medium at different temperatures. We chose a temperature range of 30-45°C, which is close to the laboratory standard (and optimal) growth temperature of 37°C.

At 30°C all strains grew with similar maximal rates, but their lag phases varied greatly (**Fig. 2A**). While the Δ*lqsS*-Δ*lqsT* and Δ*lvbR* mutant strains showed the same lag phase as the parental strain JR32, the Δ*lqsR* mutant exhibited a shortened lag phase, and the Δ*lqsA*, Δ*lqsS* and Δ*lqsT* mutants showed a prolonged lag phase. The growth initiation phenotypes of the Δ*lqsR* and Δ*lqsA* mutant strains (shortened or prolonged lag phase, respectively) were complemented at least partially by the corresponding plasmid-borne genes, and thus, are caused by the absence of these genes (**Fig. S1A)** (36). All *lqs* mutant strains as well as the *lvbR* mutant used have been complemented previously for other phenotypes, including defects in uptake by phagocytes, intracellular growth and cytotoxicity, as well as hyper-competence for DNA uptake (33, 35, 36, 39).

**FIG 2.**
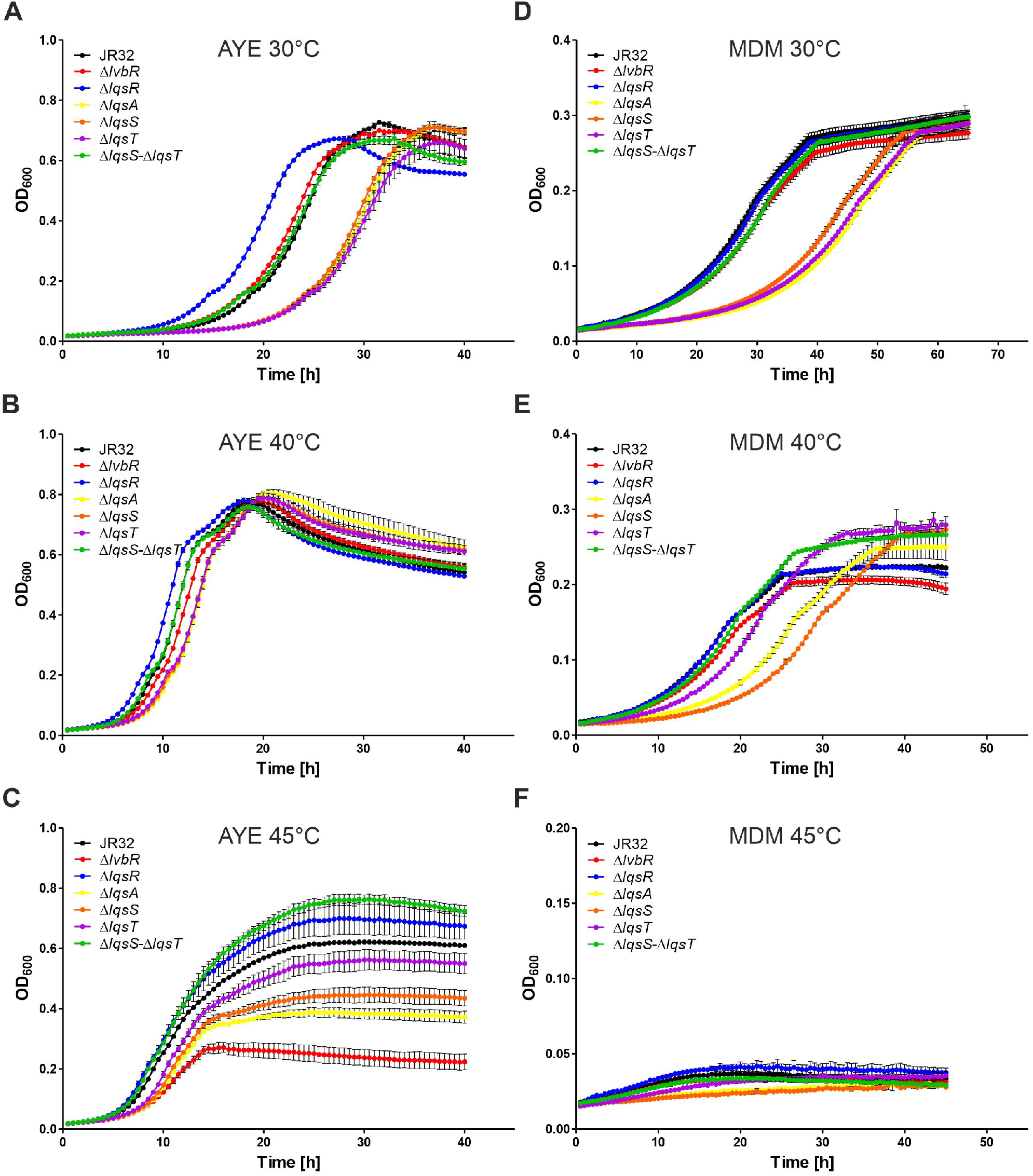
The Lqs-LvbR network regulates *L. pneumophila* growth and density in AYE medium and MDM. (A-C) *L. pneumophila* JR32, Δ*lvbR*, Δ*lqsR*, Δ*lqsA*, Δ*lqsS*, Δ*lqsT* or Δ*lqsS*-Δ*lqsT* mutant strains were grown in AYE medium for 23-24 h at 37°C and inoculated in AYE medium at an initial OD_600_ of 0.1. Growth of strains at (A) 30°C, (B) 40°C or (C) 45°C was monitored over time by measuring OD_600_ with a microplate reader. (D-F) *L. pneumophila* JR32, Δ*lvbR*, Δ*lqsR*, Δ*lqsA*, Δ*lqsS*, Δ*lqsT* or Δ*lqsS*-Δ*lqsT* mutant strains were grown in MDM for 28-29 h at 37°C and inoculated in MDM medium at an initial OD_600_ of 0.1. Growth of strains at (D) 30°C, (E) 40°C or (F) 45°C was monitored over time by measuring OD_600_ with a microplate reader. Growth curves shown are means and standard deviations of biological triplicates and representative of at least four independent experiments.

Upon growth at 40°C, the strains showed essentially the same pattern as at 30°C, except that the differences between the lag phase lengths were smaller (**Fig. 2B**). Interestingly, however, at 45°C, some strains grew with different rates and reached vastly different final cell densities (**Fig. 2C**). At 45°C, the Δ*lqsR*, Δ*lqsA*, Δ*lqsS*, Δ*lqsT* and Δ*lqsS-ΔlqsT* mutant strains grew with a similar doubling time as the parental strain JR32 (ca. 2.1-2.2 h), while the Δ*lvbR* mutant showed a longer doubling time (ca. 2.6 h), and therefore, grew significantly slower than the parental strain (**Fig. S2**). At 45°C, the Δ*lqsR* and Δ*lqsS-ΔlqsT* mutant strains reached a higher cell density compared to JR32, and the Δ*lqsA*, Δ*lqsS* and Δ*lqsT* mutant reached a lower cell density. Strikingly, the final cell density of the Δ*lvbR* mutant was less than 1/3 than that of the parental strain JR32 (**Fig. 2C**). The pronounced cell density phenotype of the Δ*lvbR* mutant strain was complemented to the full extent by the corresponding plasmid-borne gene, and thus, is caused by the absence of this gene (**Fig. S1B**). In summary, compared to the parental strain JR32, the *lqs* and the Δ*lvbR* mutant strains showed different lag phase lengths upon growth initiation in AYE medium at 30°C, and they reached different final cell densities at 45°C. In particular, the autoinducer synthase LqsA and the transcription factor LvbR positively regulate the cell density of *L. pneumophila* at higher temperatures.

### The Lqs-LvbR network regulates *L. pneumophila* growth and density in minimal defined medium

Next, we assessed the role of the Lqs-LvbR regulatory network for growth in minimal defined medium (MDM) (21). To this end, we compared growth of the parental strain JR32 with the Δ*lqsR*, Δ*lqsA*, Δ*lqsS*, Δ*lqsT*, Δ*lqsS*-Δ*lqsT* or Δ*lvbR* mutant strains in MDM at 30-45°C. At 30°C all strains grew with similar maximal rates, but – similar to growth in AYE medium – the lag phases of the Δ*lqsA*, Δ*lqsS* and Δ*lqsT* mutants were prolonged (**Fig. 2D**). Hence, the differences in the lag phases of the mutant strains are observed in the nutrient-rich and complex AYE medium, as well as in the nutrient-poorer and defined MDM. The Δ*lqsR* and Δ*lqsS*-Δ*lqsT* as well as the Δ*lvbR* mutant strains showed the same lag phase as the parental strain JR32.

Upon growth at 40°C, the strains showed a similar pattern, but only the lag phase of the Δ*lqsA* and Δ*lqsS* mutants was prolonged (**Fig. 2E**). At 45°C, growth of all strains was dramatically reduced, if not abolished (**Fig. 2F**), suggesting that compared to AYE broth, the strains cope less successfully with higher temperatures under nutrient poor conditions. Taken together, in MDM the Δ*lqsA*, Δ*lqsS* and Δ*lqsT* mutants initiated growth later than the parental strain JR32 at 30°C, and – like the other *lqs* mutants as well as the parental strain – did not grow at all at 45°C. The differences in the lag phases of the mutant strains were observed in the nutrient-rich AYE medium, as well as in the nutrient-poorer MDM and accordingly, this trait seems to be independent of the nutrients available.

### Temperature-dependent growth of environmental and clinical *Legionella* isolates

With the aim to assess the detailed growth characteristics and a role for the Lqs-LvbR system for environmental and clinical *Legionella* strains, we initially started out with a total of 19 isolates, comprising 15 *L. pneumophila* strains [serogroup (sg) 1, 5, and 6], 2 *L. longbeachae* strains and 2 *L. micdadei* strains (**Fig. S3**). Growth of these *Legionella* strains was quantitatively assessed by measuring the optical density at 600 nm (OD_600_) in AYE medium at 37°C with a 96-well microplate reader. Using this approach, we observed the following robust patterns: (i) All *L. pneumophila* strains grew with a similar doubling time (ca. 2.3-3.2 h) with the exception of the slower growing sg 6 clinical strain 526 (4.7 h), but the lag phase of the strains was different and in some cases shorter than that of the sg 1 reference strain JR32. (ii) The doubling time of the *L. longbeachae* strains (2.3 h) was similar to the *L. pneumophila* strains, but the final density was lower. (iii) The doubling time of the *L. micdadei* strains (3.3-3.6 h) was considerably higher than that of the *L. pneumophila* or *L. longbeachae* strains, and their final density was 3-4 times lower.

Based on the observed growth patterns, we picked representative strains for a detailed analysis of growth characteristics, i.e., the *L. pneumophila* environmental isolates 500 and 529 (both sg 1), 525 (sg 6) and the clinical isolate 509 (sg 1), as well as the clinical reference strains *L. pneumophila* JR32 (sg 1) and *L. longbeachae* NSW150. Like *L. pneumophila, L. micdadei* harbors the Lqs system (48); yet, due to the poor growth of the *L. micdadei* strains in AYE broth, they were not studied further. *L. longbeachae* lacks the Lqs system (49), and thus, we included this species for comparison. The selected *Legionella* strains grew with a larger doubling time and to a lower density in diluted AYE at 37°C (**Fig. S4**), indicating that the growth characteristics indeed depend on nutrient availability.

Following the growth rate of the selected *L. pneumophila* and *L. longbeachae* strains in AYE medium over a temperature range of 18°C to 50°C revealed that the growth optimum for all strains is approximately 40°C, except for strain 529, which optimally grew at 45°C (**Fig. 3A**). *L. pneumophila* strain 509 and *L. longbeachae* NSW150 were most temperature-sensitive, and at or beyond 50°C (*L. pneumophila*) or 45°C (*L. longbeachae*) the strains did not grow anymore. At the same time, the doubling times decreased from ca. 15-20 h at 18°C to ca. 2 h at 40°C (**Fig. S5A**). Intriguingly, the growth rate curve was not symmetrical: while the growth rates increased linearly in the temperature range of 18-40°C, the growth rate dropped to zero between 40-50°C (**Fig. 3A**).

**FIG 3.**
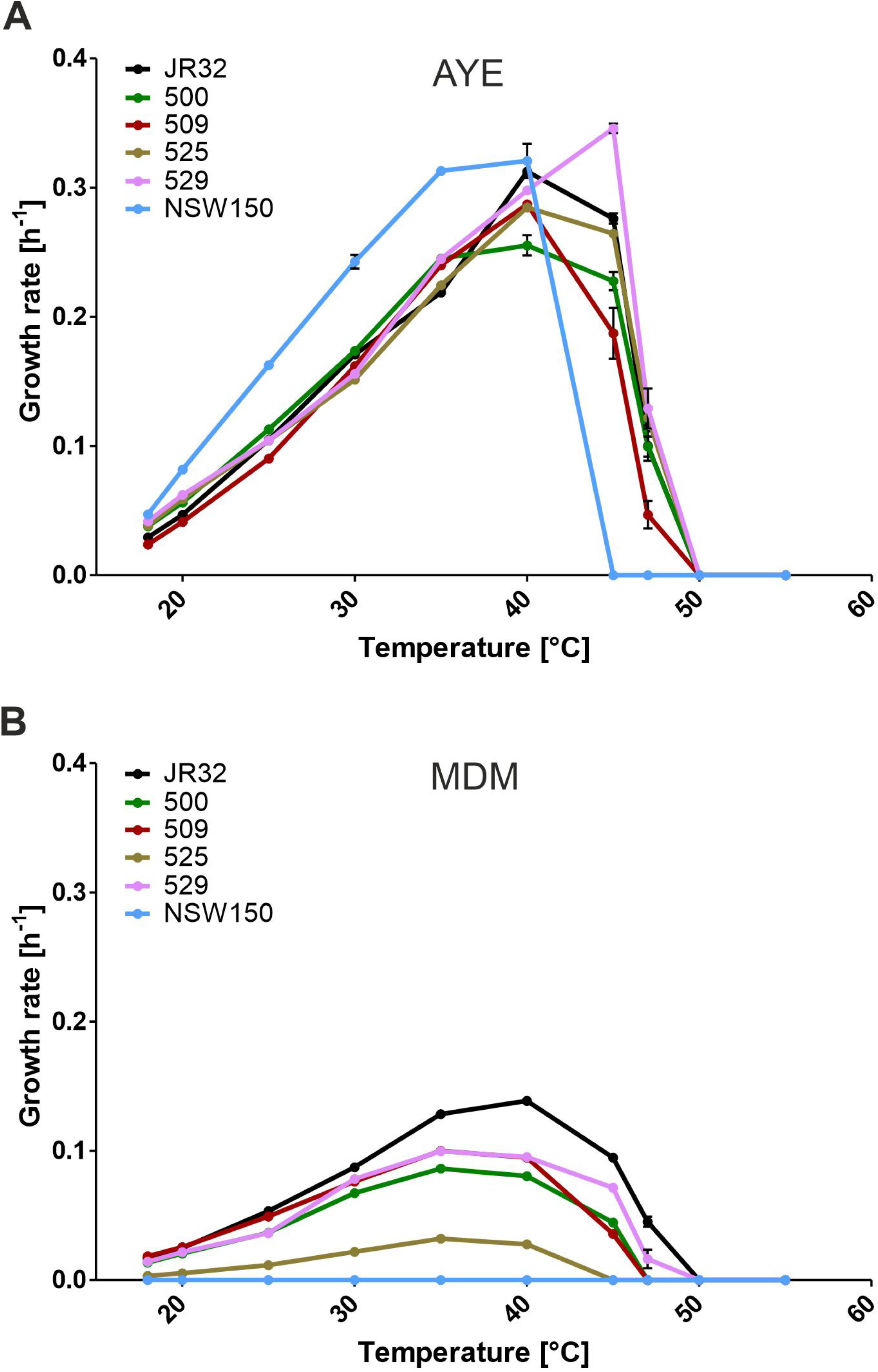
Temperature-dependent growth of environmental and clinical *Legionella* isolates. *L. pneumophila* strains JR32, 500, 509, 525, 529 and *L. longbeachae* strain NSW150 were diluted in (A) AYE medium or (B) MDM to an initial OD_600_ of 0.2 and grown at the temperatures indicated (18-55°C). Bacterial growth was determined over time by measuring OD_600_ using a microplate reader. From the resulting growth curves, the growth rates were calculated using a nonlinear regression model, only considering exponential growth phases defined in semi-logarithmic plots. Data shown are means and standard deviation of growth rates and doubling times from three independent experiments performed in biological triplicates each.

The growth rates of the *L. pneumophila* strains were ca. 3-fold lower at a given temperature in MDM, but the temperature range was similar to growth in AYE broth (**Fig. 3B**). Under these conditions, the doubling times decreased from ca. 38-52 h at 18°C to ca. 5-8 h at 40°C for most strains (**Fig. S5B**). *L. pneumophila* strain 525 (sg 6) grew exceptionally slow in MDM with a doubling time of ca. 210 h at 18°C and ca. 25 h at 40°C, and *L. longbeachae* NSW150 did not grow at all. Taken together, the doubling times of selected *L. pneumophila* and *L. longbeachae* strains decreased ca. 5-to 10-fold between growth at 18°C and the optimum of 40°C, and the doubling times of the *L. pneumophila* strains were ca. 3-fold shorter upon growth in AYE medium as compared to MDM.

### Growth onset and cell density of environmental and clinical *Legionella* strains in AYE medium

To compare growth of the selected environmental and clinical *Legionella* strains to the laboratory strain JR32 and isogenic *lqs-lvbR* mutants (**Fig. 2**), we further assessed in detail the growth characteristics at 30-45°C (**Fig. 4**). Upon growth in AYE medium at 30°C the doubling time of the 5 *L. pneumophila* strains was similar (ca. 4.0-4.6 h; **Table 1**), but the lag phase length was different (**Fig. 4A**), similar to the initially observed pattern (**Fig. S3**). While the strains 500, 525 and 529 showed a shorter lag phase compared to strain JR32, the lag phase of strain 509 was longer.

**FIG 4.**
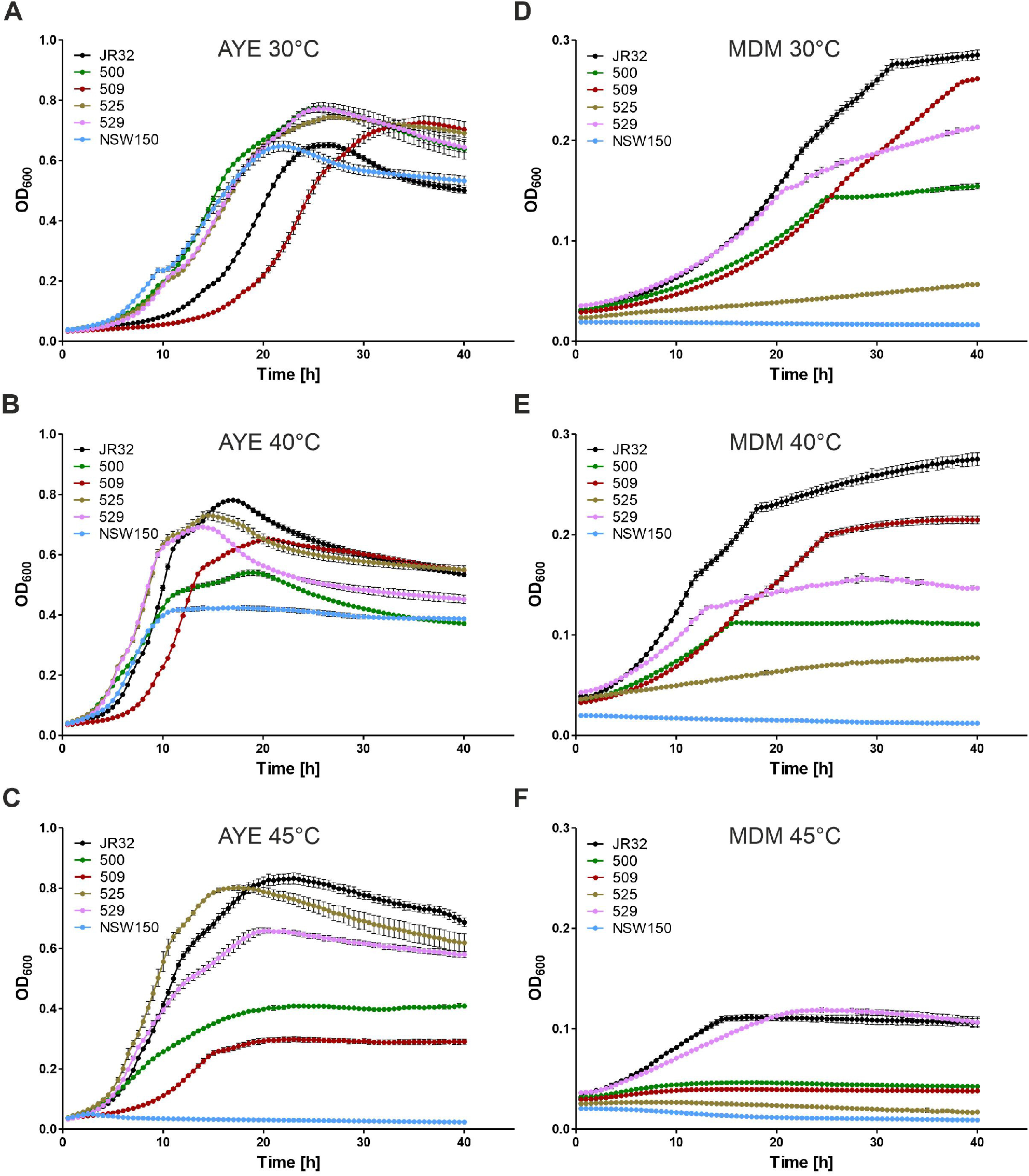
Growth onset and cell density of environmental and clinical *Legionella* strains in AYE medium and MDM. (A-C) *L. pneumophila* strains JR32, 500, 509, 525, 529 and *L. longbeachae* strain NSW150 were grown in AYE medium for 23-24 h at 37°C and inoculated in AYE medium at an initial OD_600_ of 0.2. Growth of *Legionella* strains at (A) 30°C, (B) 40°C or (C) 45°C was monitored over time by measuring OD_600_ with a microplate reader. (D-F) *L. pneumophila* strains JR32, 500, 509, 525, 529 and *L. longbeachae* strain NSW150 were grown in minimal defined medium (MDM) for 28-29 h at 37°C and inoculated in MDM at an initial OD_600_ of 0.2. Growth of strains at (D) 30°C, (E) 40°C or (F) 45°C was monitored over time by measuring OD_600_ with a microplate reader. Growth curves shown are means and standard deviations of biological triplicates and representative of six independent experiments.

**Table 1.** Doubling times of environmental and clinical *Legionella* isolates in AYE medium or MDM.

| Strain | $t_d$ 30°C<sup>a</sup> | $t_d$ 40°C<sup>a</sup> | $t_d$ 45°C<sup>a</sup> |
| --- | --- | --- | --- |
| **AYE medium** |  |  |  |
| JR32 | 4.06 ±0.03 | 2.22 ±0.02 | 2.51 ±0.04 |
| 500 | 3.99 ±0.03 | 2.72 ±0.08 | 3.05 ±0.10 |
| 509 | 4.28 ±0.02 | 2.41 ±0.01 | 3.73 ±0.37 |
| 525 | 4.58 ±0.05 | 2.44 ±0.03 | 2.62 ±0.01 |
| 529 | 4.45 ±0.03 | 2.33 ±0.01 | 2.00 ±0.02 |
| NSW150 | 2.86 ±0.06 | 2.16 ±0.09 | - |
| **MDM** |  |  |  |
| JR32 | 7.93 ±0.09 | 5.00 ±0.04 | 7.32 ±0.21 |
| 500 | 10.29 ±0.08 | 8.61 ±0.12 | 15.57 ±0.70 |
| 509 | 9.08 ±0.16 | 7.31 ±0.02 | 19.40 ±0.73 |
| 525 | 31.61 ±0.44 | 25.08 ±0.31 | - |
| 529 | 8.86 ±0.10 | 7.27 ±0.02 | 9.70 ±0.39 |
| NSW150 | - | - | - |

<sup>a</sup> Doubling times ( $t_d$ ) are indicated in [h]. Data shown are means and (±) standard deviation from doubling times derived from three independent cultures, (-) no growth observed.

Upon growth at 40°C the doubling time of the 5 *L. pneumophila* strains was reduced compared to 30°C (ca. 2.2-2.7 h; **Table 1**), the lag phase length was different, and the final cell density of *L. pneumophila* strain 500 was reduced (**Fig. 4B**). At 45°C the *L. pneumophila* strains 525, 529 and JR32 grew fastest with doubling times of 2.0-2.6 h (**Table 1**), while the *L. pneumophila* strains 500 and 509 grew slower and to much lower density (**Fig. 4C**). Intriguingly, the *L. pneumophila* strains showed a bi-phasic growth pattern, in particular at 40°C and to a somewhat lesser extent at 45°C. Finally, *L. longbeachae* NSW150 grew similarly to *L. pneumophila* (at 30°C), less densely (at 40°C) or not at all (at 45°C) (**Fig. 4A-C**, **Table 1**). Taken together, environmental and clinical *L. pneumophila* and *L. longbeachae* strains grow similarly in AYE medium after distinct lag phases at 30°C, and the strains reach different cell densities at the optimal temperature of 40°C and in particular at an elevated temperature of 45°C.

### Growth onset and cell density of environmental and clinical *Legionella* strains in minimal defined medium

Next, we assessed the growth characteristics of the 6 selected *Legionella* strains in MDM at 30-45°C. Upon growth in MDM at 30°C the *L. pneumophila* strains JR32, 509 and 529 grew with the shortest doubling time (7.9-9.1 h; **Table 1**), and strain JR32 reached the highest density (**Fig. 4D**). The *L. pneumophila* strains 500 (sg 1) and 525 (sg 6) grew much less densely, and *L. longbeachae* NSW150 did not grow at all. Compared to the nutrient-rich AYE broth, only the *L. pneumophila* strain 509 grew with a comparable pattern in MDM (log phase elongated), while the other environmental and clinical strains showed a considerably different growth pattern and also grew less densely.

Upon growth at 40°C, the growth characteristics of the strains were largely the same as at 30°C (**Fig. 4E**), but the *L. pneumophila* strains JR32, 509 and 529 grew faster with doubling times of 5.0-7.3 h (**Table 1**). At 45°C the *L. pneumophila* strains JR32 and 529 grew with doubling times similar to what was observed at 30°C, and the other strains grew barely or not at all (**Fig. 4F**). However, under these conditions, the final cell density of the strains JR32 and 529 was considerably lower, suggesting that compared to AYE broth, the strains cope less successfully with higher temperatures under nutrient poor conditions. Similar to growth in AYE medium, many *L. pneumophila* strains also showed a bi-if not tri-phasic growth in MDM, in particular the strains JR32, 500, 509 and 529 at 40°C. Taken together, the growth characteristics of the *L. pneumophila* strains JR32, 509 and 529 in MDM were qualitatively similar to AYE medium, but the final cell densities were much lower. *L. pneumophila* 525 (sg 6) barely grew, and *L. longbeachae* NSW150 did not grow at all. Since the growth pattern of same environmental and clinical strains was different in AYE broth and MDM, it appears to be dependent on the nutrients available.

### Presence and regulation of *lqs* and *lvbR* genes in environmental and clinical *Legionella* isolates

In order to possibly implicate the *lqs* and *lvbR* genes in the growth characteristics of the selected environmental and clinical *Legionella* strains, we first assessed the presence of the genes in these strains. To this end, the regions comprising P*_lqsR_*-*lqsR*, P_*lqsA*_-*lqsA*, P_*lqsS*_-*lqsS* or P_*lvbR*_-*lvbR* were amplified by PCR in the *L. pneumophila* strains JR32 (positive control), 500, 509, 525, 529 and the *L. longbeachae* strain NSW150 (negative control) (**Fig. 5A**). Using genomic DNA of these strains as a template, fragments comprising P_*lqsR*_-*lqsR* (1727 bp), P*_lqsA_*-*lqsA* (1936 bp) and P*_lqsS_*-*lqsS* (1984 bp) were amplified from all *L. pneumophila* strains tested, but not from *L. longbeachae* NSW150, in agreement with the absence of the *lqs* genes in the genome of the latter (49).

**FIG 5.**
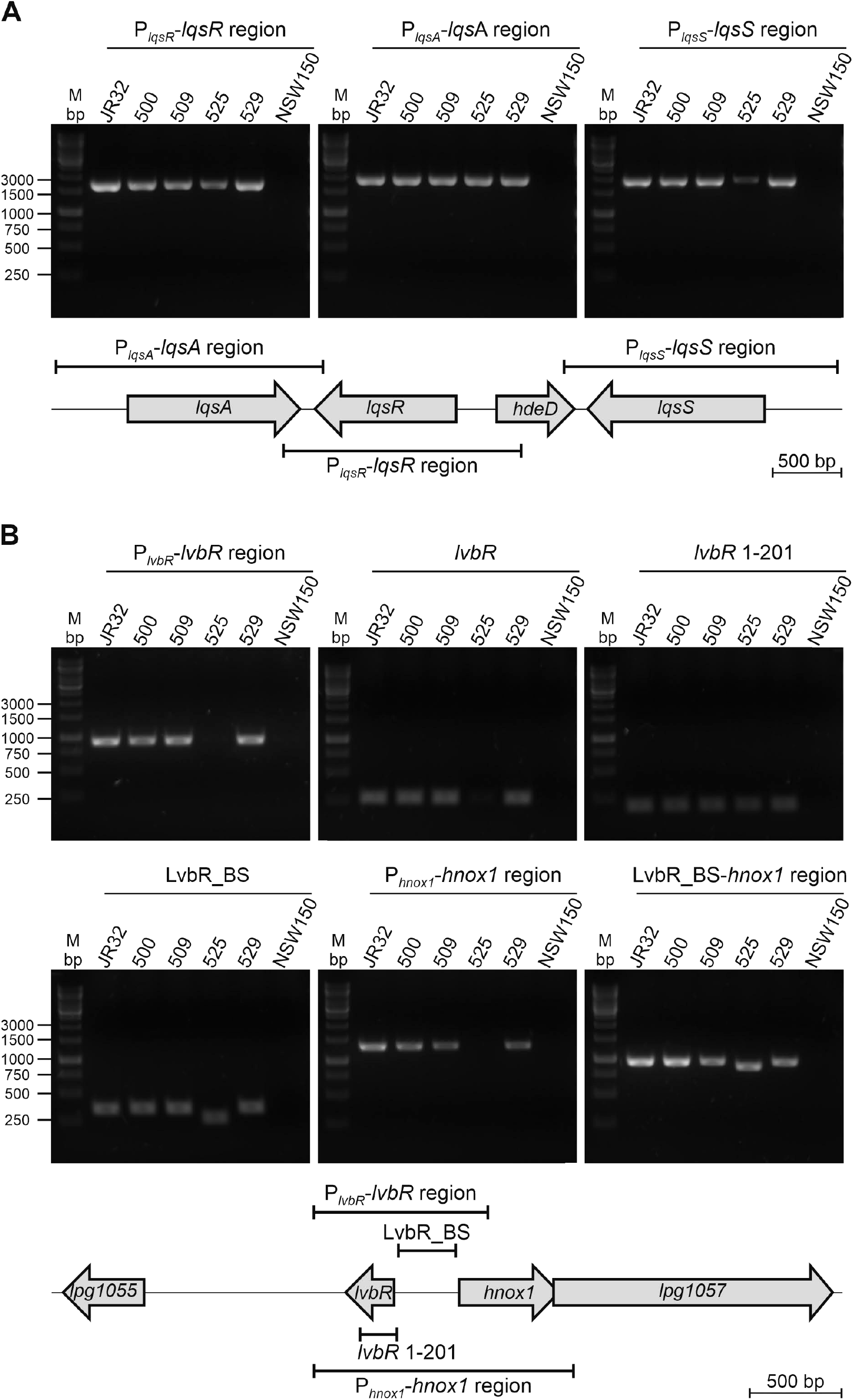
Presence of *lqsR*, *lqsA*, *lqsS*, *lvbR* and *hnox1* gene regions in environmental and clinical *Legionella* isolates. The presence of (A) P_*lqsR*_-*lqsR*, P_*lqsA*_-*lqsA*, P_*lqsS*_-*lqsS* or (B) P_*lvbR*_-*lvbR*, *lvbR*_1-201, LvbR_BS or P_*hnox1*_-*hnox1* in the genomes of *L. pneumophila* strains JR32 (positive control), 500, 509, 525, 529 or *L. longbeachae* strain NSW150 (negative control) was assessed by PCR amplification of the corresponding genes (and putative promoters) and/or intergenic regions indicated. PCR products were analyzed by agarose gel electrophoresis and visualized with MaestroSafe staining: P_*lqsR*_-*lqsR* (1727 bp), P_*lqsA*_-*lqsA* (1936 bp), P_*lqsS*_-*lqsS* (1984 bp), and P_*lvbR*_-*lvbR* (948 bp), *lvbR* (264 bp), *lvbR*_1-201 (201 bp), LvbR_BS (347 bp), P_*hnox1*_-*hnox1* (1344 bp) and LvbR_BS-*hnox1* (969 bp). Schemes depicting the *lqs* or the *lvbR*-*hnox1* genomic region and the PCR fragments amplified are shown below the gels.

The fragments P*_lvbR_*-*lvbR* (948 bp), *lvbR* (264 bp) and P_*hnox1*_-*hnox1* (1344 bp) were amplified in the *L. pneumophila* strains JR32, 500, 509, and 529, but neither in *L. pneumophila* 525 nor in *L. longbeachae* NSW150 (**Fig. 5B**). However, the fragments *lvbR*_1-201 (201 bp), LvbR_BS (LvbR binding site (39); 347 bp) and LvbR_BS-*hnox1* (969 bp) were amplified in all *L. pneumophila* strains tested including strain 525, indicating that the *lvbR* as well as the *lvbR*-*hnox1* intergenic region and *hnox1* are indeed present in the genome of the latter strain. Strain 525 harbors an LvbR_BS region, which is shorter than in the other strains tested, and an *lvbR* gene, which appears to be a more distantly related homologue. Sequencing of the LvbR_BS intergenic region and part of the *lvbR* and *hnox1* genes of the strains JR32, 500, 509, 525 and 529 revealed the presence of almost identical *lvbR* and *hnox1* genes and an LvbR_BS intergenic region of strain 525, which is indeed 80 nucleotides shorter (**Fig. S6**). *L. longbeachae* NSW150 likely does not harbor an *lvbR* homologue, and the most closely related protein is a putative YdnC-like transcription factor (29% identity of 55/178 amino acids; Evalue 7 × e^-4^). Taken together, these results revealed that the *lqsR*, *lqsA*, *lqsS* and *lvbR* genes are present in all *L. pneumophila* strains analyzed in detail in this study.

Next, we tested the regulation of the *lqs* and *lvbR* genes in environmental and clinical *L. pneumophila* isolates. The genome sequences for these strains are not available, and therefore, we used the corresponding promoters of strain JR32. Accordingly, we transformed the strains JR32, 500, 509, 525 and 529 with reporter constructs comprising transcriptional fusions of different promoters from strain JR32 and *gfp* (P*_lqsR_*-*gfp*, P*_lqsA_*-*gfp*, P_*lqsS*_-*gfp*, P*_lvbR_*-*gfp* or P*_hnox1_*-*gfp*) and quantified GFP production by fluorescence (**Fig. 6**). This approach revealed that all environmental and clinical *L. pneumophila* isolates tested expressed P*_lqsR_*-*gfp* (**Fig. 6A**), P*_lqsA_*-*gfp* (**Fig. 6B**), P*_lqsS_*-*gfp* (**Fig. 6C**), P_*lvbR*_-*gfp* (**Fig. 6D**) and P*^hnox1^*-*gfp* (**Fig. 6E**). The expression pattern of P*_lqsR_*, P*_lqsS_* and P*_lvbR_*, as well as of P*_lqsA_* and P_*hnox1*_, were similar in the environmental and clinical *L. pneumophila* isolates. Interestingly, some environmental and clinical *L. pneumophila* isolates expressed P_*lvbR*_ with high intensity, as opposed to the parental strain JR32, which did not express P*_lvbR_* (**Fig. 6D**), as observed previously (39). Since all of the environmental and clinical strains harbor *lqsS* (**Fig. 5A**), and therefore presumably produce the negative regulator of LvbR, LqsS (35, 39), another mechanism likely accounts for the observation that the strains indeed express P_*lvbR*_.

**FIG 6.**
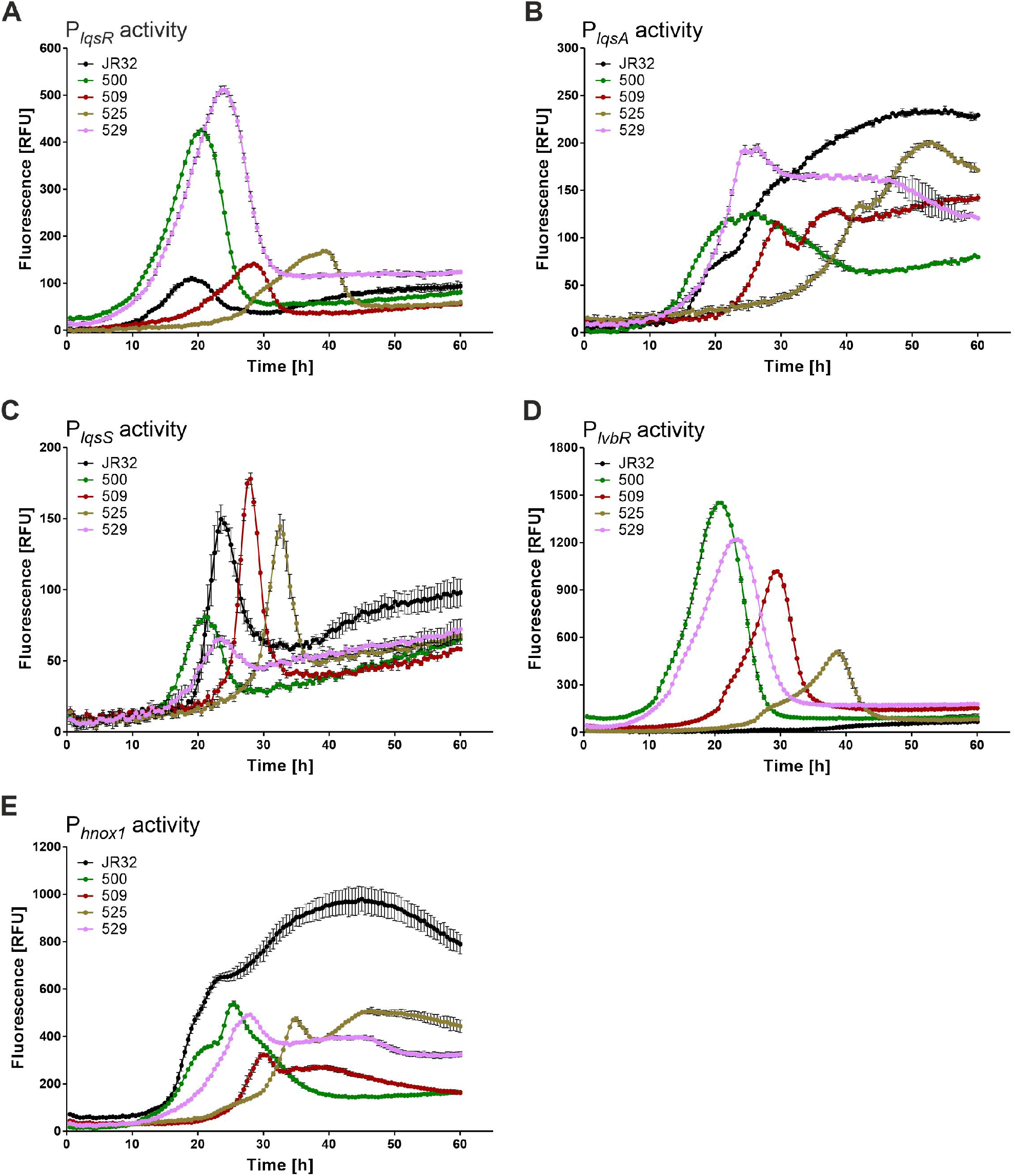
Regulation of *lqsR*, *lqsA*, *lqsS*, *lvbR* and *hnox1* promoters in environmental and clinical *Legionella* isolates. *L. pneumophila* strains JR32, 500, 509, 525 and 529 harboring (A) P_*lqsR*_-*gfp* (pRH037), (B) P_*lqsA*_-*gfp* (pRH038), (C) P_*lqsS*_-*gfp* (pRH051), (D) P_*lvbR*_-*gfp* (pRH023) or (E) P_*hnox1*_-*gfp* (pRH026) reporter constructs (promoters of strain JR32) were grown in AYE medium for 23-24 h at 37°C and inoculated in AYE medium at an initial OD_600_ of 0.2. Strains were incubated at 30°C, and GFP fluorescence was measured over time (relative fluorescence units, RFU) using a microplate reader. Data shown are means and standard deviations of biological triplicates and representative of three independent experiments.

The expression of P_*lqsR*_, P_*lqsS*_ and P_*lvbR*_ in the environmental and clinical *L. pneumophila* isolates peaked at different times (**Fig. 6**), which precisely correspond to the different lag phase lengths of the strains, and accordingly, to their transition from the exponential to the stationary growth phase (**Fig. 7**). On the other hand, P*_lqsA_* and P*_hnox1_* showed a similar, but more complex expression profile, which was not exclusively linked to the transition from the exponential to the stationary growth phase (**Fig. 7**).

**FIG. 7.**
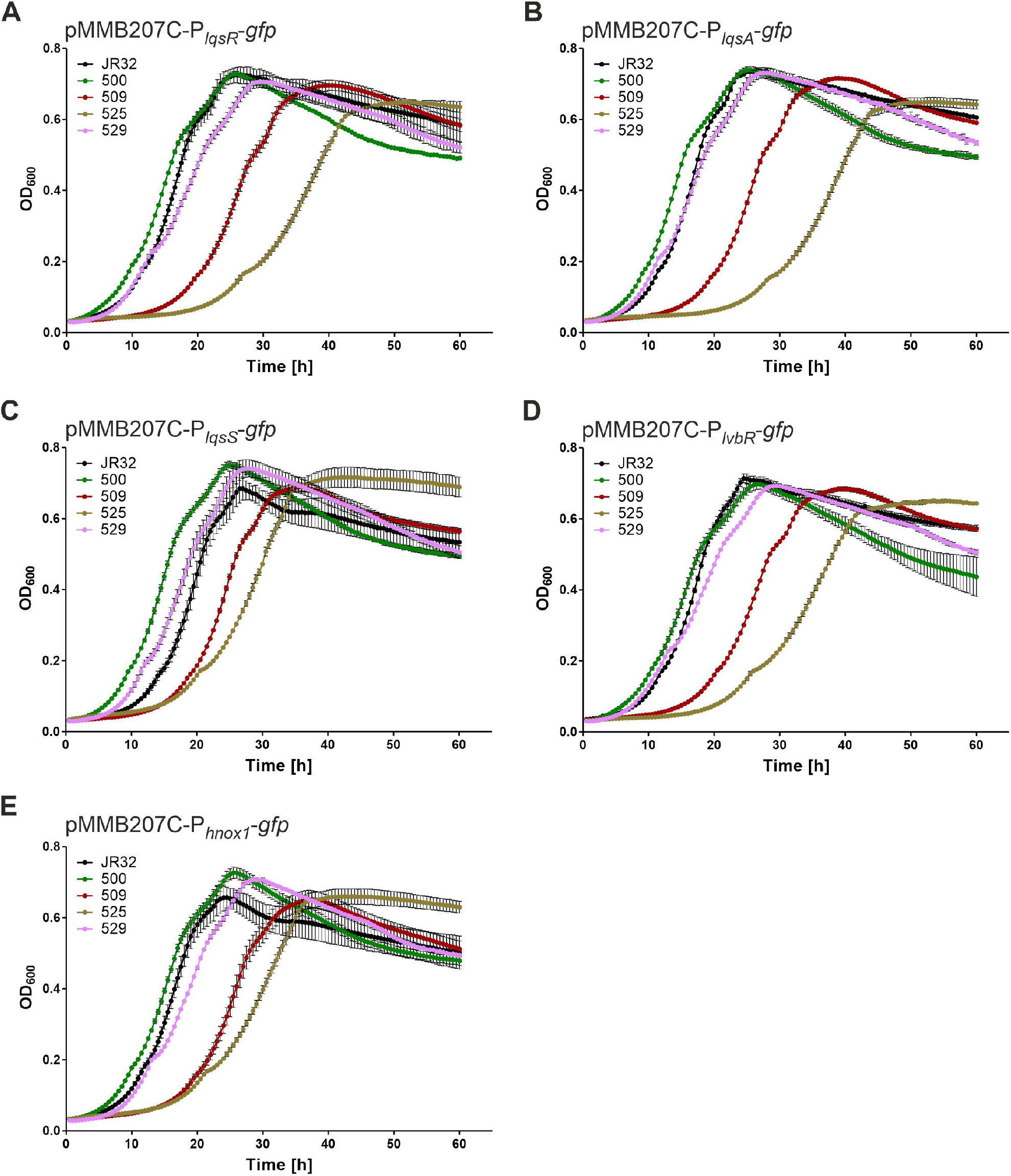
Growth of environmental and clinical *L. pneumophila* isolates harboring promoter reporters. *L. pneumophila* strains JR32, 500, 509, 525 and 529 containing (A) P_*lqsR*_-*gfp* (pRH037), (B) P_*lqSA*_-*gfp* (pRH038), (C) P_*lqsS*_-*gfp* (pRH051), (D) P_*lvbR*_-*gfp* (pRH023) or (E) P_*hnox1*_-*gfp* (pRH026) reporter constructs (promoters of strain JR32) were grown in AYE medium for 23-24 h at 37°C and inoculated in AYE medium at an initial OD_600_ of 0.2. Bacterial growth at 30°C was followed over time by measuring OD_600_ with a microplate reader. Data shown are means and standard deviations of biological triplicates and representative of three independent experiments.

The *L. pneumophila* strains JR32, 500, 509, 525, and 529 harboring the empty plasmid pMMB207C-RBS showed very low fluorescence (**Fig. S7A**). The growth profile of these strains at 30°C (**Fig. S7B**) was very similar to the growth profiles of the strains harboring no plasmid (**Fig. 4A**), but distinct from the growth profiles of strain 525 harboring one of the promoter reporters producing GFP (**Fig. 7A-E**). Therefore, GFP production rather than the presence of the plasmid itself seems to account for the delayed growth of strain 525, but not the other *L. pneumophila* strains tested. In summary, all environmental and clinical *L. pneumophila* isolates tested expressed P_*lqsR*_, P_*lqsS*_ and P*_lvbR_* of strain JR32 with a single peak in the post-exponential growth phase, and the strains expressed P*_lqsA_* and P*_hnox1_* of strain JR32 with a more complex expression profile. These results suggest that the Lqs-LvbR network is active in the environmental and clinical *L. pneumophila* strains.

## Discussion

In this study we analyzed in detail the growth characteristics and the role of the Lqs-LvbR network for temperature-dependent growth rate and cell density of a *L. pneumophila* reference and mutant strains, as well as of select environmental and clinical *L. pneumophila* and *L. longbeachae* isolates. For the *L. pneumophila* reference strain, we implicated components of the Lqs system in growth onset, doubling time and cell density (**Fig. 2**). Hence, the Lqs system not only regulates *L. pneumophila* virulence, motility, competence, and a genomic fitness island, but also growth characteristics of planktonic bacteria (**Fig. 1**). Compared to the parental strain JR32, a Δ*lqsR* mutant strain lacking the response regulator LqsR showed a reduced lag phase at 30°C in AYE medium and reached a higher cell density at 45°C (**Fig. 2**). The Δ*lqsA*, Δ*lqsS* and Δ*lqsT* mutants lacking the autoinducer synthase LqsA or one of the homologous sensor kinases, LqsS or LqsT, respectively, exhibited a longer lag phase and reached only a lower cell density. Intriguingly, the components of the Lqs system regulated growth resumption of sessile *L. pneumophila* in a similar manner: compared to the parental strain JR32, the percentage of Δ*lqsA*, Δ*lqsS* or Δ*lqsT* mutant strains resuming growth was lower, while the percentage of Δ*lqsR* and the Δ*lqsS*-Δ*lqsT* double mutant was higher (46).

Intriguingly, growth resumption and final cell density of the *lqs* and *lvbR* mutant strains is not correlated to their virulence and transmission phenotypes. Both the Δ*lqsR* and the Δ*lvbR* mutant strains are severely impaired for virulence (36, 39), yet growth resumption characteristics and final cell density at the temperatures tested are different (**Fig. 2**). Analogously, the Δ*lqsA* mutant does not show a severe virulence phenotype compared to the parental strain JR32 (35), yet growth resumption and final cell density of the two strains are distinct (**Fig. 2**). Finally, the Δ*lqsS*, Δ*lqsT* and Δ*lqsS*-Δ*lqsT* double mutant are all impaired for virulence (33, 35), yet the individual sensor kinase mutants reached a lower cell density than the parental strain at 45°C, while the Δ*lqsS*-Δ*lqsT* double mutant reached a higher cell density. A possible explanation for the latter observation is that LqsS and LqsT regulate many genes in a complex and reciprocal manner (33), and hence, the Δ*lqsS*-Δ*lqsT* double mutant might cancel out or revert phenotypes of the single mutants. In any case, the effects of *L. pneumophila* Lqs-LvbR components on planktonic growth resumption and final cell density are apparently not directly linked to the virulence phenotypes of the corresponding mutant strains.

The Lqs system and the α-hydroxyketone autoinducer LAI-1 are not strictly conserved among *Legionella* species, but α-hydroxyketone-mediated signaling and cell-cell communication seem to be widely distributed among environmental bacteria (27). A recent bioinformatics analysis for the presence of the *lqs* cluster in 58 *Legionella* genomes revealed three categories of species (48): 19 harbored a complete *lqs* cluster, 20 did not possess *lqsA* but maintained the receptor *lqsR* and/or the sensor kinase *lqsS*, and 19 did not have any of the *lqs* genes. These results are in agreement with the notion that the Lqs system functions in intra-as well as inter-species communication. In agreement with a broad distribution of the Lqs-LvbR network are the findings that components are not only present in the reference strain JR32, but also in the genomes of the clinical and environmental *L. pneumophila* strains analyzed in detail here (**Fig. 5**). Interestingly, the P_*lqsR*_, P_*lqsA*_ and P_*lqsS*_ promoters from strain JR32 are expressed to a similar if not higher degree in the clinical and environmental *L. pneumophila* strains, and P_*lvbR*_ is highly expressed in these strains, but not in JR32 (**Fig. 6**), confirming previous findings (39).

In addition to the Lqs system, we found that the pleiotropic transcription factor LvbR plays a major role for doubling time and cell density of *L. pneumophila* (**Fig. 2**). In AYE medium, the Δ*lvbR* mutant strain resumed growth like the parental strain at 30°C (**Fig. 2A**), but exhibited a dramatically reduced cell density at 45°C (**Fig. 2C**). In MDM, the Δ*lvbR* mutant strain behaved similarly to the parental strain JR32 (**Figs. 2D-F**). Hence, preferentially under nutrient rich conditions and at elevated temperatures, the LvbR transcription factor functions as an important regulator of *L. pneumophila* cell density. The low final cell density reached by the Δ*lvbR* mutant strain in AYE medium might be reflected in the different transcriptomes of the mutant compared to the parental strain JR32 (39). An inspection of genes differentially produced in the Δ*lvbR* mutant versus the parental strain revealed that many flagellar genes are up-regulated, while genes implicated in the metabolism of amino acids, lipids and carbohydrates are indeed down-regulated.

In the laboratory strain JR32, LvbR is negatively regulated by LqsS (35, 39) (**Fig. 6D**). All environmental and clinical strains tested harbor *lqsS* (**Fig. 5A**) as well as *lvbR* (**Fig. 5B**, **S6B**). Based on expression studies with the corresponding promoters of strain JR32, *lqsS* as well as *lvbR* are expressed in the environmental and clinical strains (**Fig. 6CD**). Therefore, *lvbR* does not seem to be negatively regulated by *lqsS* in these strains, and the production of LvbR might not be linked to the Lqs system.

In *L. pneumophila* strain JR32, LvbR links the Lqs system to c-di-GMP signaling (39, 40), and therefore, it is likely that the second messenger c-di-GMP controls temperature-dependent growth rate and cell density of the pathogen. *L. pneumophila* JR32 harbors 22 genes predicted to encode proteins with domains implicated in c-di-GMP synthesis, hydrolysis and recognition (50, 51). However, only a few components of the c-di-GMP regulatory network have been characterized, including the Hnox1-Lpg1057 system, comprising the diguanylate cyclase inhibitor Hnox1 and the GGDEF/EAL domain-containing diguanylate cyclase Lpg1057 (41, 42) (**Fig. 1**). It is currently not known, which components of the c-di-GMP regulatory network play a role in the temperature-dependent control of *L. pneumophila* growth rate and cell density under given conditions.

Recently, we analyzed the growth of sessile *L. pneumophila* on surfaces and in biofilm using single cell techniques (46). We found that sessile *L. pneumophila* exhibits phenotypic heterogeneity and adopts growing and non-growing (“dormant”) states in biofilms and microcolonies formed in AYE medium. The establishment of phenotypic heterogeneity was controlled by the temperature, the Lqs system and LvbR. The Lqs system and LvbR were found to determine the ratio between growing and non-growing (“persister”) sessile subpopulations, as well as the frequency of growth resumption (“resuscitation”) and microcolony formation of individual bacteria. The microcolony growth rate was the same for the parental strain JR32 and *lqs* mutant strains, similar to what we observed for planktonic growth in AYE medium at 30°C or 40°C (**Figs. 2A, B**). LvbR regulated the growth resumption of sessile *L. pneumophila* (46), but not the lag phase length of planktonic *L. pneumophila* growing at 30°C or 40°C (**Figs. 2A, B**), and therefore, substantial differences between the LvbR-dependent growth regulation of sessile and planktonic bacteria exist.

Other regulatory systems implicated in the survival of *L. pneumophila* in aqueous environments are the “stringent response” and the alternative sigma factor RpoS (σ^S^/σ^38^) (52), as well as the two-component system (TCS) LetAS (53). The stringent response is triggered upon production of the second messenger guanosine 3’,5’-bispyrophosphate (ppGpp) by the RelA and SpoT synthases (54–56), which in turn are activated by amino acid starvation (57) and inhibition of fatty acid biosynthesis (56). In response to ppGpp the stationary phase sigma factor RpoS controls the expression of *L. pneumophila* virulence and transmission (54, 58), and the LetAS TCS also regulates these traits (28, 59–61). RpoS and to a lesser extent LetA control the production of LqsR in stationary growth phase (36), and thus, the stringent response is linked to the Lqs system. For planktonic growth of *L. pneumophila* in broth, minimal media and aqueous environments this implicates that nutrient starvation likely upregulates the Lqs system, and hence, promotes density-dependent regulation.

In addition to the role of the intrinsic Lqs-LvbR system for growth onset, doubling time and cell density of *L. pneumophila*, we also assessed the role of the extrinsic (environmental) cues temperature and nutrient availability for growth of *Legionella* spp. (**Figs. 3, 4, S3-5**). Compared to *L. pneumophila*, *L. longbeachae* appeared to be more susceptible to elevated temperatures in AYE medium (**Fig. 3, 4**). While *L. pneumophila* strains grew up to 47°C and ceased to grow at 50°C, *L. longbeachae* reached only a lower final cell density upon growth at 40°C and ceased to grow already at 45°C (**Fig. 3, 4**). The most temperature-sensitive *L. pneumophila* strains were the environmental isolate 500 and the clinical isolate 509 (both sg 1), which at a growth temperature of 45°C reached only a 2-3-fold lower cell density in AYE medium compared to the reference strain JR32 (**Fig. 4C**). The final cell density of these strains did not correlate with the differences in the lag phase length, since compared to JR32 strain 500 or 509 showed a shortened or extended lag phase, respectively, at 30°C. In general, the temperature-dependent differences in the final cell density became even more pronounced in MDM. Under these conditions, only strain JR32 and the environmental isolate 529 grew, albeit to only a low density, while all other *L. pneumophila* strains no longer grew (**Fig. 4D-F**). The environmental and clinical *L. pneumophila* strains all harbored the *lqsR*, *lqsA*, *lqsS* and *lvbR* genes in their genomes, and thus, the Lqs-LvbR network might play a role for the temperature-dependent growth patterns, but not for the differences observed among the strains. Accordingly, the reason(s) for the vast differences in growth resumption and final density of natural *Legionella* isolates in response to temperature are unknown.

Overall, the results documented in this study are in agreement with a general role for quorum sensing (Lqs) and c-di-GMP signaling (LvbR) for growth onset, doubling time and final cell density of *L. pneumophila* under different conditions. Future studies will identify the “downstream” components implicated in the temperature-, Lqs- and LvbR-dependent regulation of *L. pneumophila* growth under different extra- and intracellular conditions.

## Material and Methods

### Growth of *Legionella* strains

The *Legionella* strains used in this study are listed in **Table 2**. *Legionella* strains were grown on CYE agar plates (62) and in liquid cultures with *N*-(2-acetamido)-2-aminoethanesulfonic acid (ACES)-buffered yeast extract (AYE) medium (63) for 23-24 h or in minimal defined medium (MDM) (21) for 28-29 h at 37°C on a wheel (80 rpm). To obtain growth curves, *Legionella* strains were diluted from cultures grown in AYE medium to an initial OD_600_ of 0.1 or 0.2 with AYE medium or MDM into 96-well microplates (100 μl/well, polystyrene, 353072, Falcon) and incubated at the temperatures indicated in the figure legends while orbitally shaking. Bacterial growth was monitored over time in triplicates by measuring the optical density at 600 nm (OD_600_) using a microplate reader (Synergy H1 or Cytation5 Hybrid Multi-Mode Reader, BioTek). *L. pneumophila* strains harboring empty control plasmids (pTS10 or pNT28) or complementation plasmids (pAK18, pTS02, pTS04) were cultivated and analyzed in AYE medium supplemented with chloramphenicol (Cm; 5 μg/ml).

**Table 2.**
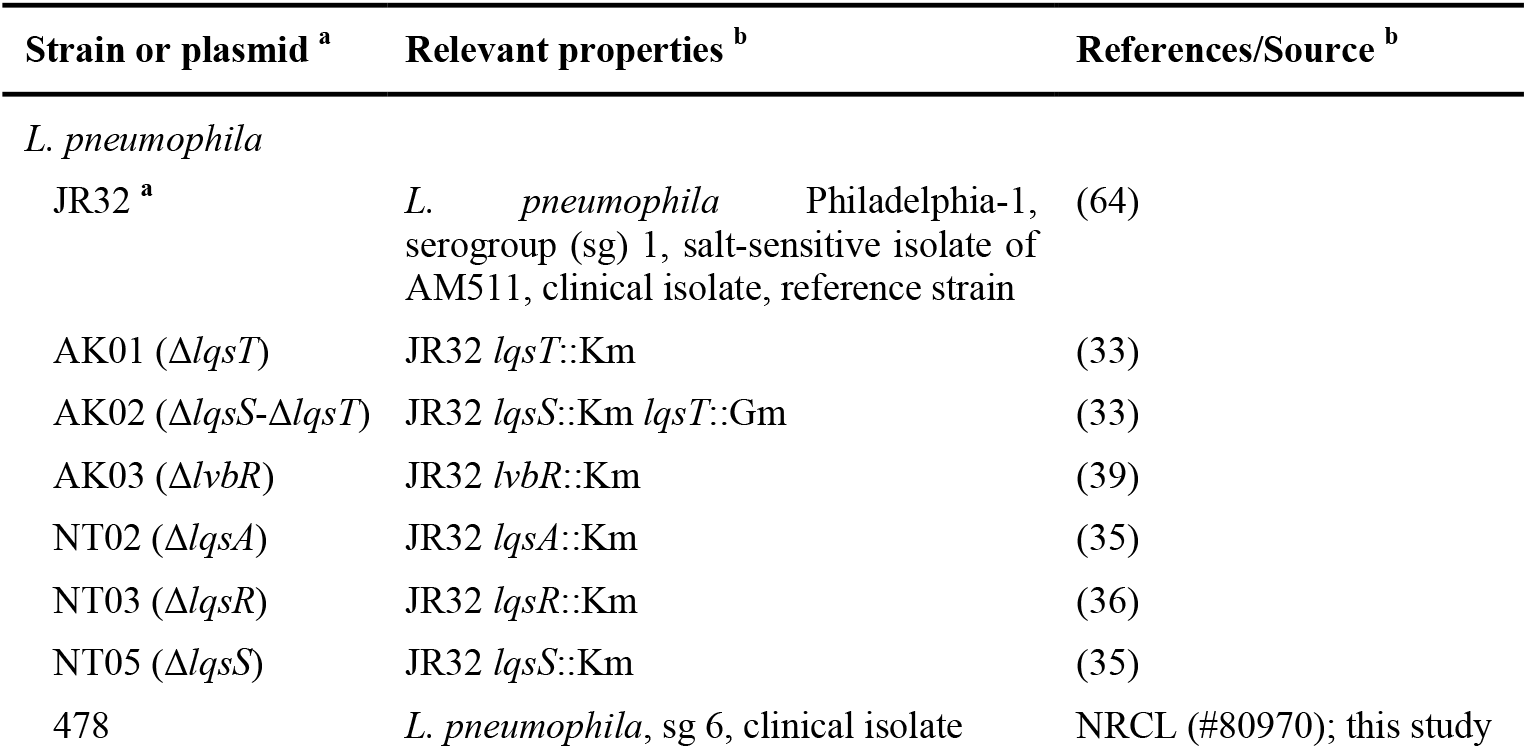
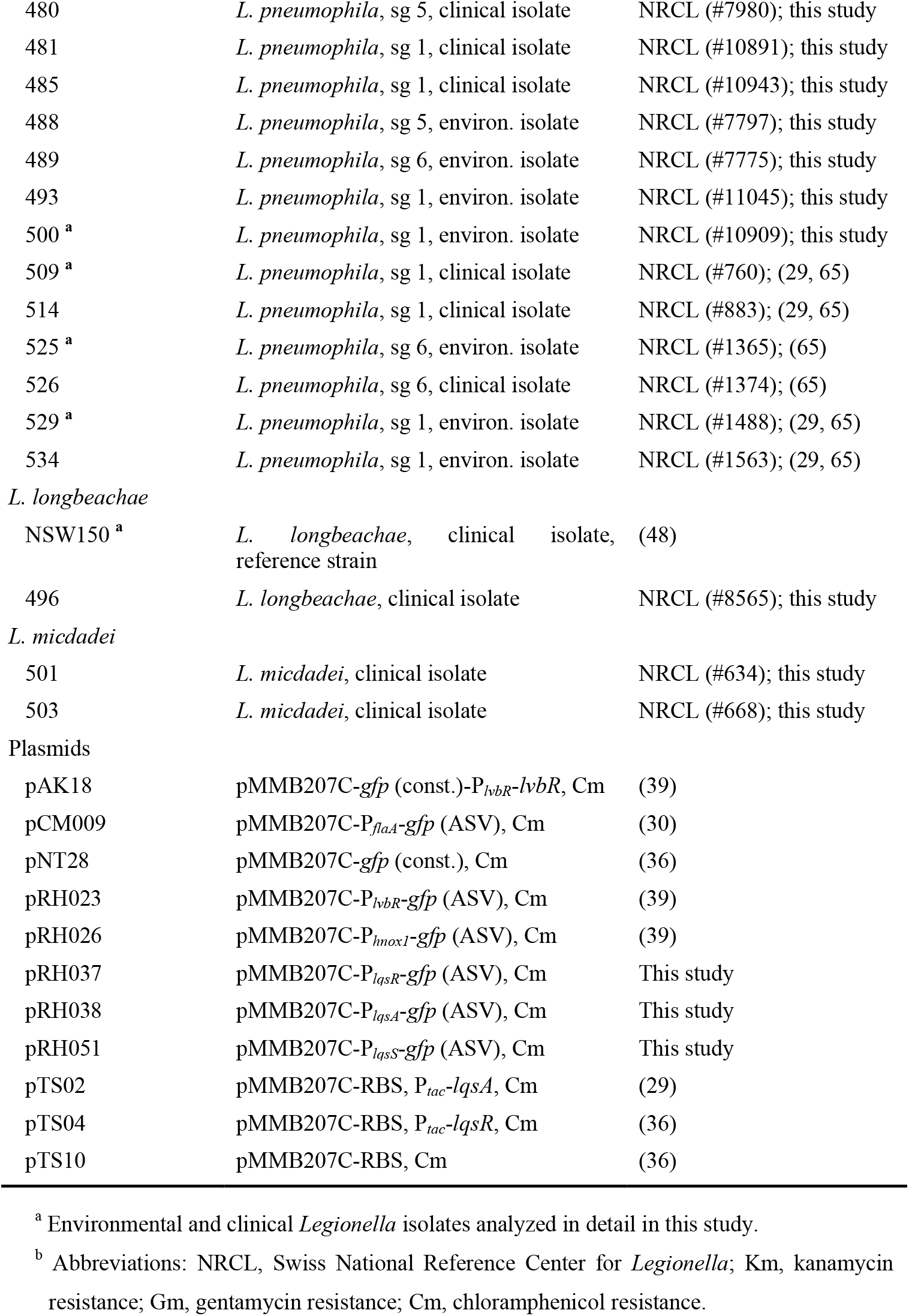
*Legionella* strains and plasmids used in this study.

The experiments and the statistics were performed in biological triplicates, as defined by individual bacterial cultures growing in separate wells of a microtiter plate. The biological triplicates were repeated at least twice in independent experiments, as defined as bacterial cultures originating from different pre-cultures, which yielded practically identical results. In most cases, the experiments were independently repeated 3-5 times. The resulting growth curves were analyzed with the software GraphPad Prism (Version 5.04). Growth rates (maximum) and doubling times (minimum) were calculated using a curve fit with a nonlinear regression model, considering only the fastest exponential growth as defined in semi-logarithmic plots.

### Plasmid construction

Cloning was performed according to standard protocols, and the primers used for PCR reactions are listed in **Table 3**. Plasmids were amplified and isolated from *E. coli* TOP10 (Invitrogen) using commercially available kits (Macherey-Nagel). Reporter plasmids pRH037, pRH038 and pRH051 containing a transcriptional P*_lqsR_*-, P*_lqsA_*-, or P*_lqsS_*-*gfp* (ASV) fusion were constructed by amplifying the insert by PCR using the primer pair oRH188/189, oRH190/191 or oRH223/224, respectively, and genomic JR32 DNA as a template. PCR products were cloned with the NEBuilder HiFi DNA Assembly reaction (NEB) into the *Sac*I and *Xba*I sites of pCM009 (30), thereby replacing P_*flaA*_. The resulting constructs were verified by DNA sequencing.

**Table 3.**
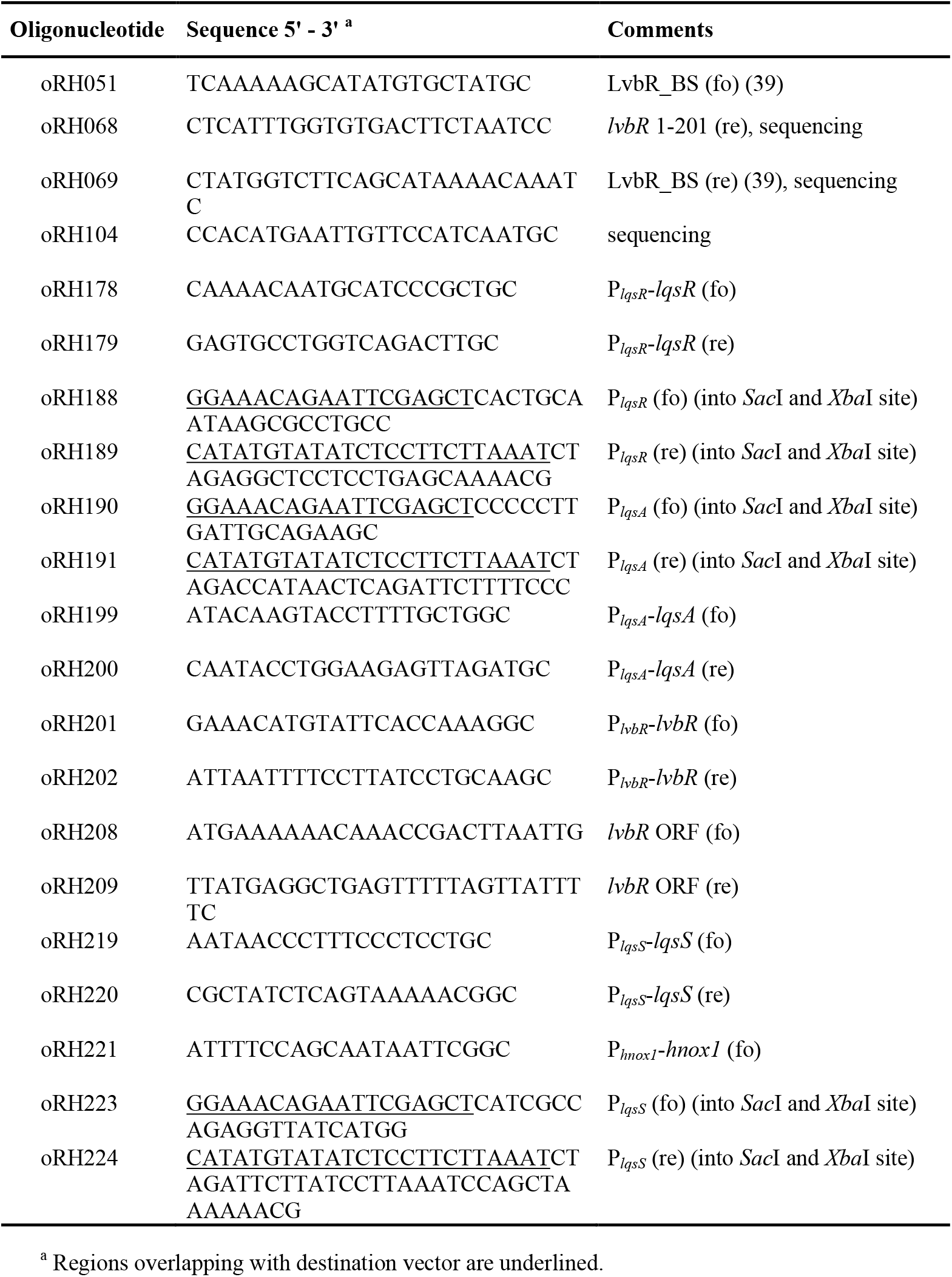
Oligonucleotides used in this study.

### Genomic DNA extraction, PCR amplification and sequencing

Genomic DNA was extracted from *Legionella* strains cultured for 18 h in AYE medium using the GenElute^™^ Bacterial Genomic DNA Kit (NA2100, Sigma-Aldrich) and used as template for PCR reactions. Genes and putative promoter regions were amplified by standard PCR protocols using the primer pairs oRH178/179 (P_*lqsR*_-*lqsR*), oRH199/200 (P_*lqsA*_-*lqsA*), oRH219/220 (P_*lqsS*_-*lqsS*), or oRH201/202 (P*_lvbR_*-*lvbR*), oRH208/209 (*lvbR*), oRH208/068 (*lvbR*_1-201), oRH051/069 (LvbR_BS), oRH221/202 (P_*hnox1*_-*hnox1*) or oRH221/051 (LvbR_BS-*hnox1*), and the PCR fragments were subsequently analyzed by agarose gel electrophoresis. DNA bands were visualized with MaestroSafe staining (Labgene Scientific SA) and imaged using a FluorChem SP imager (Alpha Innotech). For sequence analysis, the *lvbR-hnox1* gene regions were amplified from genomic DNA by PCR using primer pair oRH104/068. PCR products were analyzed by agarose gel electrophoresis, purified (gel and PCR clean-up kit, Macherey-Nagel) and subsequently sequenced by the use of primers oRH068, 069 or 104 (Sanger sequencing performed by Microsynth). An alignment was created with the resulting sequences by the program CLC Main Workbench 7.

### GFP reporter assays

*L. pneumophila* JR32, 500, 509, 525 or 529 strains harboring P_*lqsR*_-, P*_lqsA_*-, P*_lqsS_*-, P*_lvbR_*- or P_*hnox1*_-*gfp* (ASV) fusion reporter plasmids (pRH037, pRH038, pRH051, pRH023 or pRH026) or empty pMMB207C plasmids (pTS10) were grown for 23-24 h at 37°C in AYE medium supplemented with chloramphenicol (Cm; 5 μg/ml). Strains were subsequently inoculated from these cultures at an initial OD_600_ of 0.2 in AYE medium/Cm into a black clear bottom 96-well plate (100 μl/well, polystyrene, 353219, Falcon) and incubated at 30°C while orbitally shaking. GFP production and bacterial growth were monitored in triplicates by measuring fluorescence (excitation, 485 nm; emission, 528 nm; gain, 50) and the OD_600_ with a microplate reader (Synergy H1 or Cytation5 Hybrid Multi-Mode Reader, BioTek). Blank value (AYE medium) was subtracted from all values, and numbers are expressed as relative fluorescence units (RFU) or OD_600_.

## Abbreviations

Icm/Dot: intracellular multiplication/defective organelle trafficking
LAI-1: *Legionella* autoinducer-1
LCV: *Legionella*-containing vacuole
Lqs: *Legionella* quorum sensing
LvbR: *Legionella* virulence and biofilm regulator
c-di-GMP: cyclic di-guanosine monophosphate
T4SS: type IV secretion system

## Acknowledgements

We would like to thank Barbara Borer and Arnd Gildemeister (Geberit International AG) for stimulating discussions. The project was funded by Geberit International AG. Work in the group of H.H. was supported by the Swiss National Science Foundation (31003A_175557, 310030_200706), a research grant from the University of Zürich awarded to R.H., and the Institute of Medical Microbiology.

The funders had no role in study design, data collection and analysis, decision to publish, or preparation of the manuscript. The authors declare no conflict of interest.

## Notes

### Competing Interest Statement

The authors have declared no competing interest.

### Summary of Updates

Supplemental Information provided.

