## Supplemental Figures S1-S7 for "The *Legionella* Lqs-LvbR regulatory network controls temperature-dependent growth onset and bacterial cell density"

**Running title:** Growth regulation by the Lqs-LvbR regulatory network

**Key words:** amoeba, bacterial physiology, biofilm, cell-cell communication, *Legionella*, metabolism, transcription factor, quorum sensing.

**Abbreviations:** Icm/Dot, intracellular multiplication/defective organelle trafficking; LAI-1, *Legionella* autoinducer-1; LCV, *Legionella*-containing vacuole; Lqs, *Legionella* quorum sensing; LvbR, *Legionella* virulence and biofilm regulator, c-di-GMP, cyclic di-guanosine monophosphate; T4SS, type IV secretion system.

### Supplementary Figures

#### Figure S1

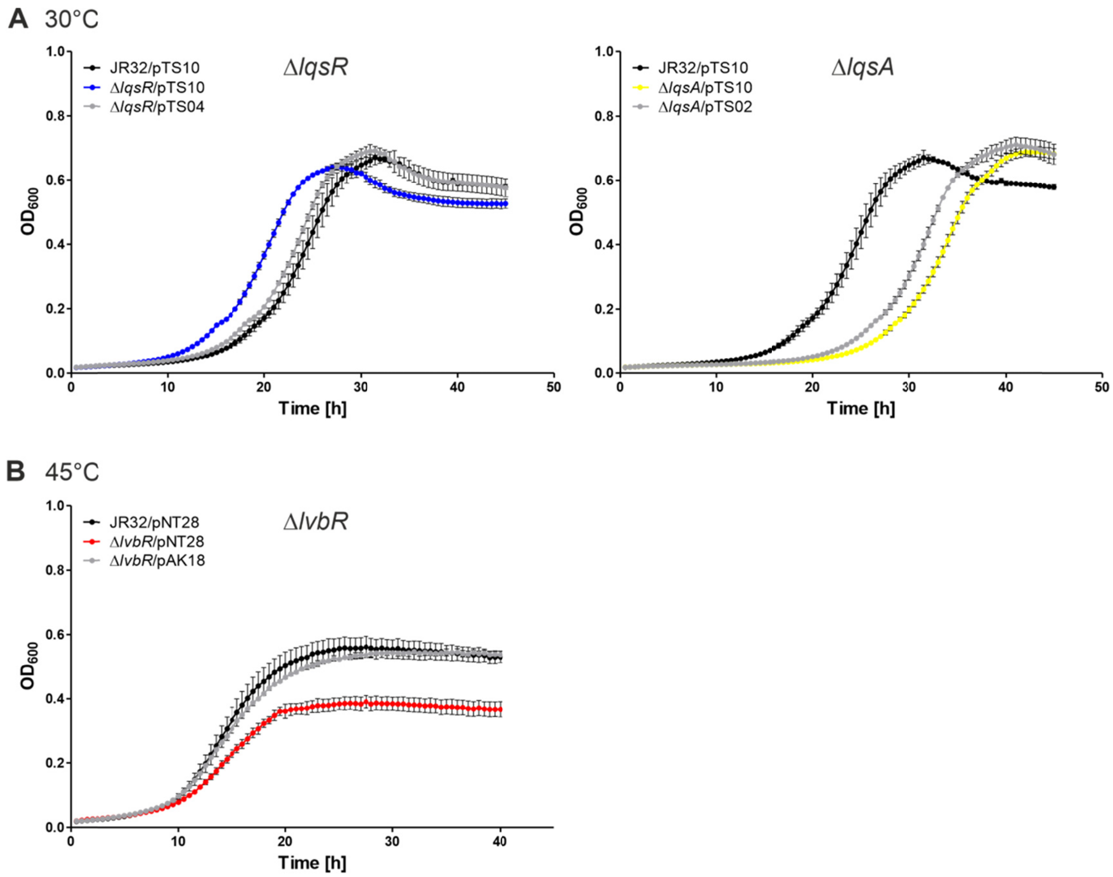


**FIG S1. Complementation of *L. pneumophila* Δ*lqsR*, Δ*lqsA* and Δ*lvbR* growth phenotypes.** *L. pneumophila* JR32 and (**A**) Δ*lqsR* and Δ*lqsA* or (**B**) Δ*lvbR* mutant strains harboring an empty plasmid (pTS10 or pNT28) or a complementation plasmid expressing either *lqsR* ($P_{tac}$, pTS04), *lqsA* ($P_{tac}$, pTS02) or *lvbR* ($P_{lvbR}$, pAK18) were grown in AYE medium for 23-24 h at 37°C. The strains were inoculated at an initial $OD_{600}$ of 0.1, and growth at (**A**) 30°C or (**B**) 45°C was monitored over time by measuring $OD_{600}$ with a microplate reader. Growth curves shown are means and standard deviations of biological triplicates.

**Figure S2**

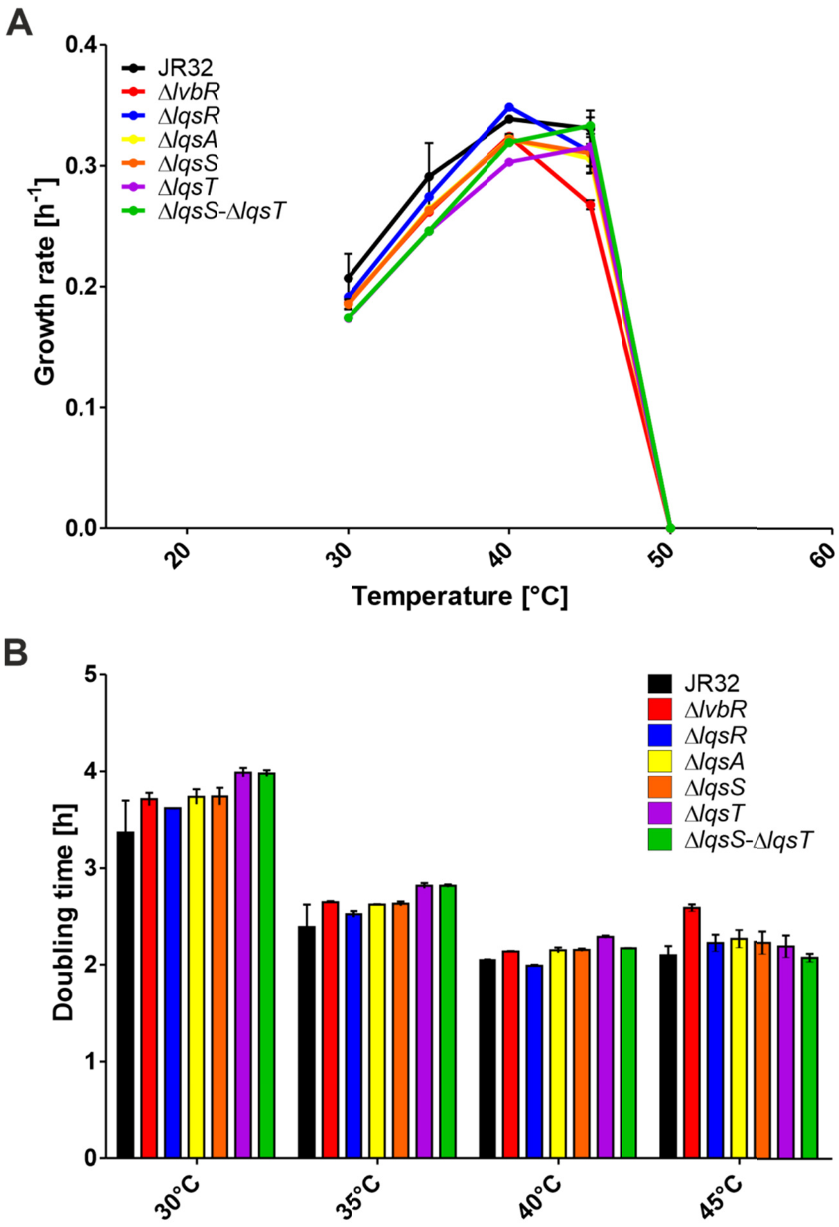


**FIG S2. Temperature-dependent growth of *L. pneumophila lqs* and *lvbR* mutant strains.** *L. pneumophila* JR32, Δ*lqsR*, Δ*lqsA*, Δ*lqsS*, Δ*lqsT*, Δ*lqsS*-Δ*lqsT* or Δ*lvbR* mutant strains were grown in AYE medium for 23-24 h at 37°C and inoculated in AYE medium at an initial $OD_{600}$ of 0.1. The bacterial growth at the temperatures indicated (30-50°C) was monitored over time by measuring $OD_{600}$ with a microplate reader. (**A**) Growth rates and (**B**) doubling times were calculated from resulting growth curves using a nonlinear regression model, only considering exponential growth phases defined in semi-logarithmic plots. Data shown are means and standard deviation of growth rates and doubling times from two independent experiments performed in biological triplicates each.

**Figure S3**

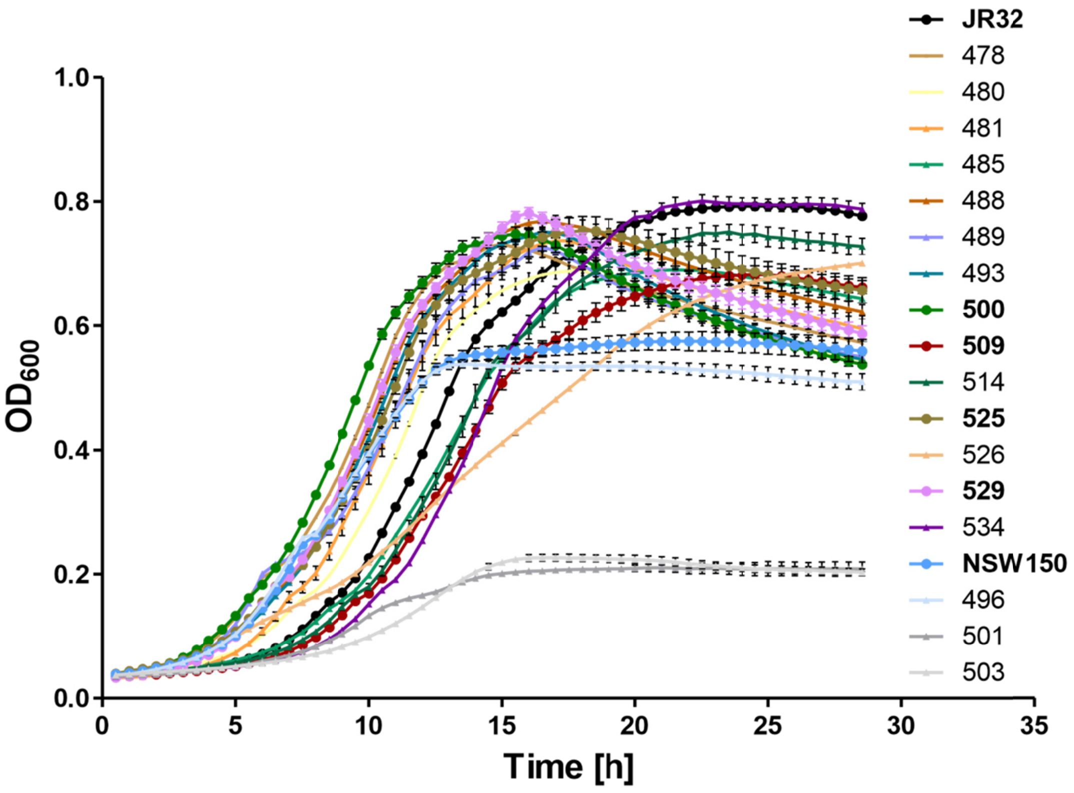


**FIG S3. Growth curves of environmental and clinical *Legionella* isolates in AYE medium.** *L. pneumophila* (JR32, 478, 480, 481, 485, 488, 489, 493, 500, 509, 514, 525, 526, 529, 534), *L. longbeachae* (NSW150, 496) and *L. micdadei* (501, 503) strains were grown in AYE medium for 23-24 h at 37°C and inoculated in AYE medium at an initial $OD_{600}$ of 0.2. The strains were incubated at 37°C, and growth was followed by measuring $OD_{600}$ over time with a microplate reader. Growth curves shown are means and standard deviations of biological triplicates and representative of at least two independent experiments. Strains selected for detailed analysis are depicted with circle symbols and indicated in bold.

**Figure S4**

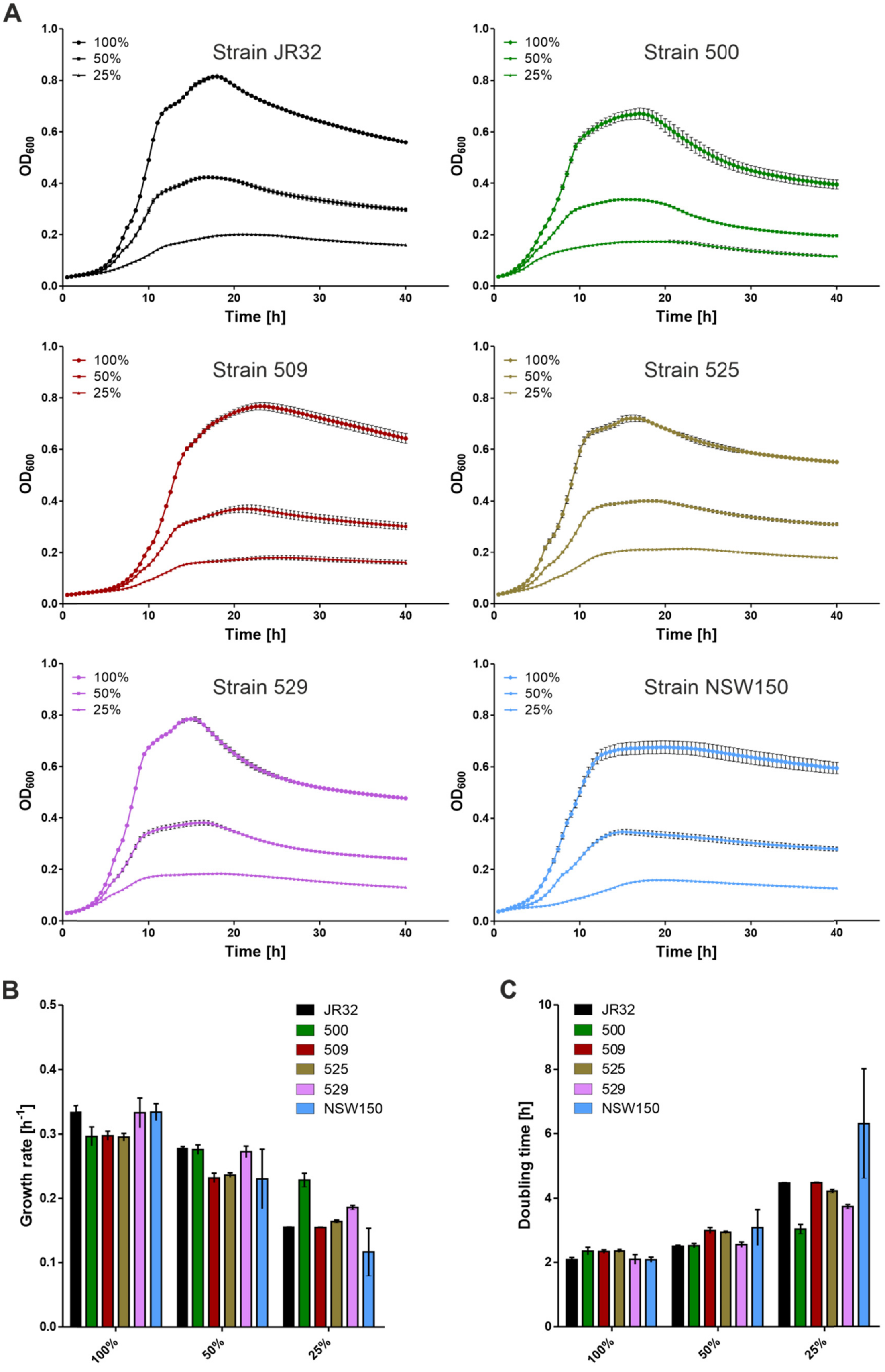
A
Strain JR32
Strain 500
Strain 509
Strain 525
Strain 529
Strain NSW150
100%
50%
25%
$OD_{600}$
Time [h]
B
Growth rate [h$^{-1}$]
C
Doubling time [h]
JR32
500
509
525
529
NSW150
100%
50%
25%

**FIG S4 (overleaf). Growth curves of environmental and clinical *Legionella* isolates in diluted AYE medium.** *L. pneumophila* strains JR32, 500, 509, 525, 529 and *L. longbeachae* strain NSW150 were grown in AYE medium for 23-24 h at 37°C and inoculated in 25%, 50% or 100% AYE medium at an initial $OD_{600}$ of 0.2. (**A**) Bacterial growth was monitored over time at 40°C by measuring $OD_{600}$ with a microplate reader. From the resulting growth curves (**B**) the growth rates and (**C**) the doubling times were calculated using a nonlinear regression model, only considering exponential growth phases defined in semi-logarithmic plots. Growth curves shown are means and standard deviations of biological duplicates and representative of two independent experiments.

**Figure S5**

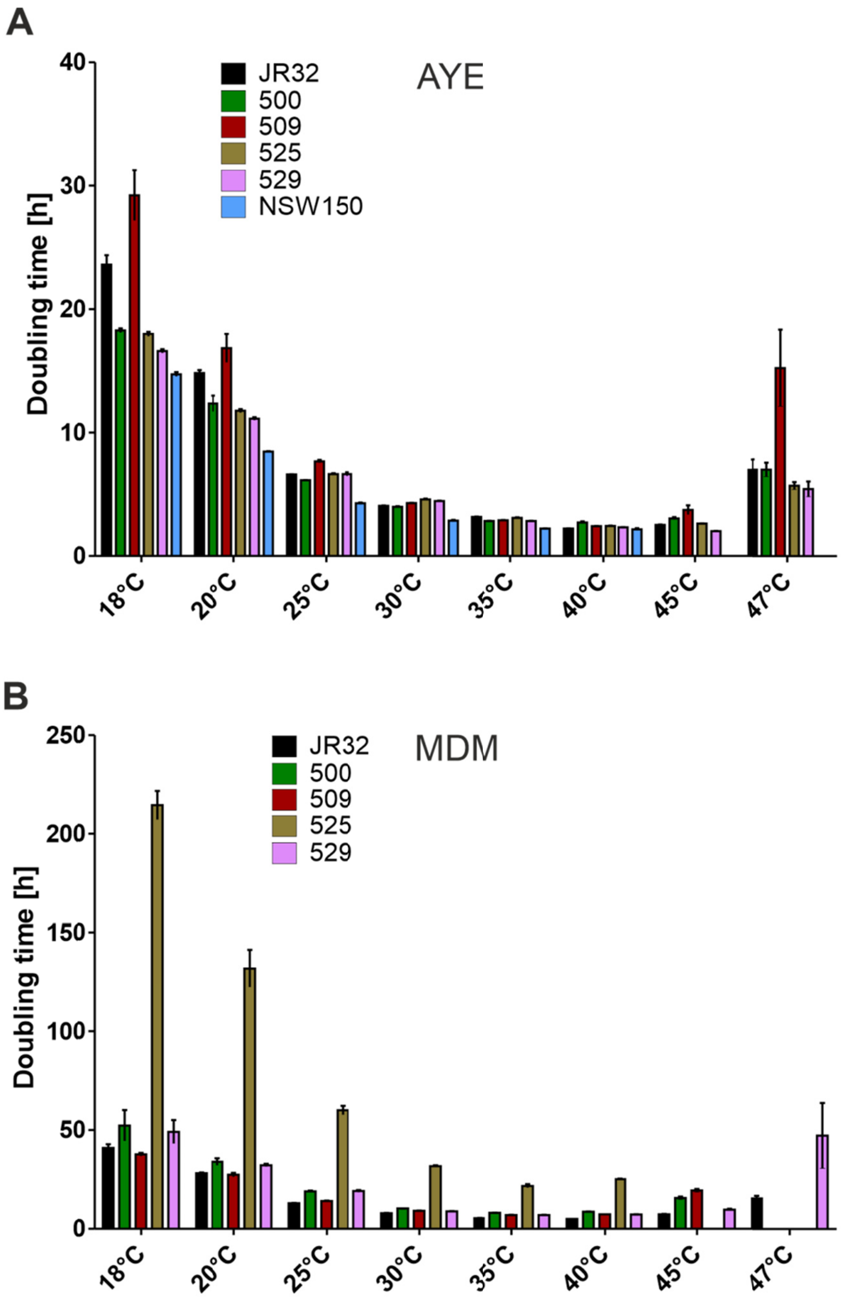


**FIG S5. Temperature-dependent growth of environmental and clinical *Legionella* isolates.** *L. pneumophila* strains JR32, 500, 509, 525, 529 and *L. longbeachae* strain NSW150 were diluted in (**A**) AYE medium or (**B**) MDM to an initial $OD_{600}$ of 0.2 and grown at the temperatures indicated (18-55°C). The growth was determined over time by measuring $OD_{600}$ using a microplate reader. The doubling times were calculated from the growth curves using a nonlinear regression model, only considering exponential growth phases defined in semi-logarithmic plots. Data shown are means and standard deviation of doubling times derived from three independent experiments analyzed in biological triplicates each.

**Figure S6**

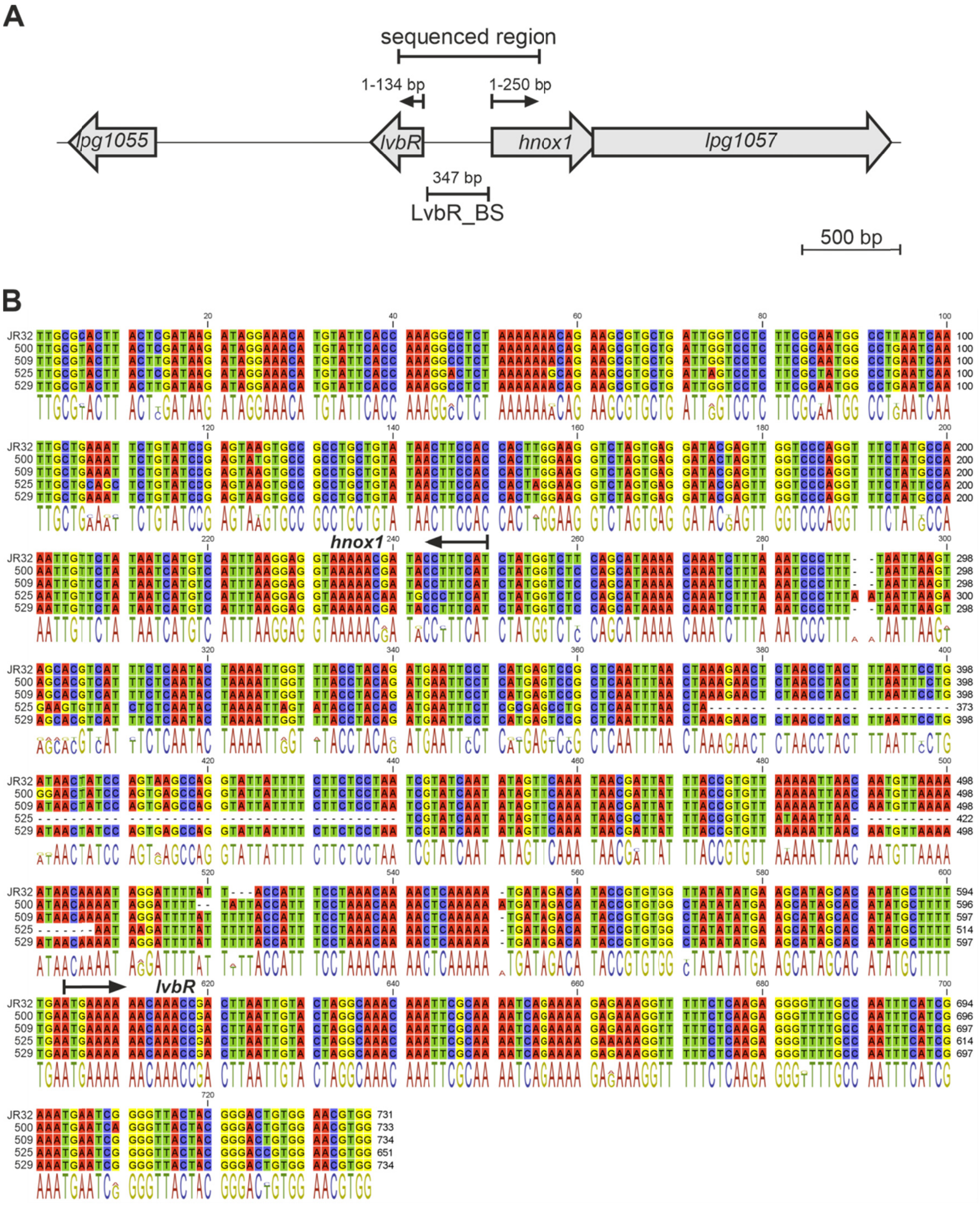


**FIG S6. *LvbR-hnox1* gene region of environmental and clinical *L. pneumophila* isolates.** (**A**) Scheme of the *L. pneumophila lvbR*-*hnox1* gene region. The "sequenced region", including 134 bp of *lvbR*, LvbR_BS and 250 bp of *hnox1*, was amplified by PCR from genomic DNA of the *L. pneumophila* strains JR32, 500, 509, 525 and 529, and subsequently sequenced. (**B**) Alignment of the resulting nucleotide sequences. The open reading frames of the *lvbR* or *hnox1* genes are indicated by arrows.

**Figure S7**

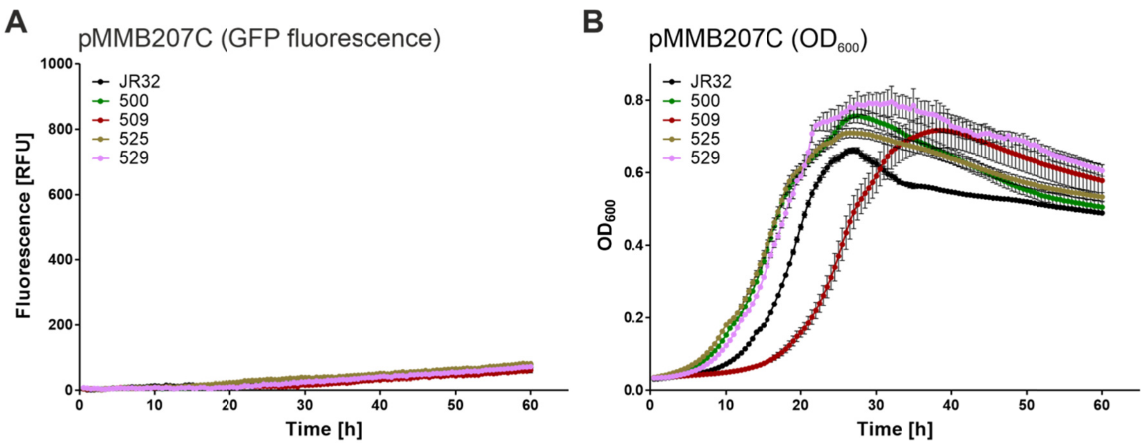


**FIG S7. Fluorescence and growth of *L. pneumophila* strains harboring plasmid pMMB207C-RBS.** The *L. pneumophila* strains JR32, 500, 509, 525 and 529 containing the empty vector pMMB207C-RBS (pTS10) were grown in AYE medium for 23-24 h at 37°C and inoculated in AYE medium at an initial $OD_{600}$ of 0.2. The strains were incubated at 30°C, and (**A**) GFP fluorescence (relative fluorescence units, RFU) and (**B**) growth ($OD_{600}$) was measured over time using a microplate reader. Data shown are means and standard deviations of biological triplicates and representative of two independent experiments.
